# A rapid, multiplex digital PCR assay for *EGFR*, *KRAS*, *BRAF*, *ERBB2* variants and *ALK*, *RET*, *ROS1*, *NTRK1* gene fusions in non-small cell lung cancer

**DOI:** 10.1101/2023.03.09.531949

**Authors:** Bryan Leatham, Katie McNall, Hari K. Subramanian, Lucien Jacky, John Alvarado, Dominic Yurk, Mimi Wang, Donald C. Green, Gregory J. Tsongalis, Aditya Rajagopal, Jerrod J. Schwartz

## Abstract

Digital PCR (dPCR) is emerging as an ideal platform for the detection and tracking of genomic variants in cancer due to its high sensitivity and simple workflow. The growing number of clinically-actionable cancer biomarkers creates a need for fast, accessible methods that allow for dense information content and high accuracy. Here, we describe a proof-of-concept amplitude modulation based multiplex dPCR assay capable of detecting 12 single nucleotide and indel variants in *EGFR*, *KRAS*, *BRAF*, and *ERBB2,* 14 gene fusions in *ALK, RET, ROS1, NTRK1,* and *MET* exon 14 skipping present in non-small cell lung cancer (NSCLC). We also demonstrate the use of multi-spectral target signal encoding to improve the specificity of variant detection by reducing background noise up to 11-fold. The assay reported an overall 100% PPA and 98.5% NPA compared to a sequencing-based assay in a cohort of 62 human FFPE samples. In addition, the dPCR assay rescued actionable information in 10 samples that failed to sequence, highlighting the utility of a multiplexed digital assay as a potential reflex solution for challenging NSCLC samples.

## 1 Introduction

Lung cancer is the leading cause of cancer death in the United States, with a projected 350 deaths per day in 2022 [1]. Fortunately, there are a growing number of advancements in screening and treatment response monitoring, as well as targeted therapies and immunotherapies, that have improved clinical management for patients with advanced NSCLC [1, 2]. For example, there are now over a dozen different precision medicines targeting driver genes and network pathways [3]. Despite these improvements in treatment options for NSCLC patients, there remain significant challenges with current molecular test options that critically limit treating patients with the right drugs. Constraints including test accessibility, sample availability, and the lack of consistent payor reimbursement for diagnostic tests have prevented widespread utilization of precision medicines [4]. Challenges such as insufficient or poor quality samples, and slow turnaround time [5], have further hindered broad adoption. For example, in a 2022 multisource database investigation, nearly 50% of patients were unable to benefit from precision medicines due to factors linked with obtaining biomarker results; 18% received inaccurate results due to test limitations or errors; and 4% started on a less precise treatment due to prolonged test turnaround time [6]. Therefore, there is an outstanding need for rapid, comprehensive, reliable, and low-cost methods that can identify patients as eligible for precision treatment and clinical trials.

Massively parallel, or next-generation sequencing (NGS), is the leading approach to profile both primary tumor samples and peripheral cell-free nucleic acids for clinically-actionable biomarkers. A main advantage of this method is that sequence information of entire genes and regions of the genome is generated, which enables comprehensive detection of variants present. However, there are also key challenges with sequencing-based approaches, including: test failures due to insufficient specimen volume, nucleic acid isolation yields, or failed library preparation [7], complex and time-consuming laboratory workflows and bioinformatics analysis [8], and high instrumentation and reagent cost [9, 10]. These factors have limited both the successful processing of clinical samples and the types of institutions performing these assays. dPCR is an emerging alternative to NGS for cancer biomarker testing due to its simple workflow, low sample input requirements, high sensitivity, fast turnaround time, and low cost [11, 12, 13]. However, the clinical utility of conventional dPCR remains limited due to its inherent multiplexing limitation to assess all actionable biomarkers in a single assay with a limited amount of sample. To overcome this, several methods have been proposed to increase digital PCR information content through amplification curve analysis [14, 15], melt curve analysis [14, 15], and amplitude modulation [16]. However, none of these methods have yet been developed into a comprehensive assay that generates a complete set of actionable information because of complexities in workflows.

Here we describe a proof of concept TaqMan<u>®</u>-based amplitude modulation-based digital PCR panel [HDPCR, see 17] for multiplexed detection of relevant variants seen in NSCLC, including 12 single nucleotide or insertion/deletion DNA variants, 14 RNA fusion variants, and *MET* exon 14 skipping (Table S1). All DNA variants and RNA fusion variants detected by this panel were selected based on NCCN guideline recommendations and the association of targeted therapies for advanced or metastatic NSCLC [18]. The amplitude modulation scheme relies on standard, low cost, TaqMan probe hydrolysis that is concentration limited to deterministically program unique fluorescent signatures for each analyte. Given that modern PCR instruments incorporate photodetectors with a wide dynamic range, multiple targets each with a corresponding unique fluorescent intensity can be multiplexed within one channel. The panel also leverages multi-spectral signal encoding for some analytes to create a form of error detection code [19] that improves the specificity of analyte detection beyond standard TaqMan PCR by lowering the effective background noise. Together, the digital PCR panel enables a three-hour turn-around-time of results from isolated nucleic acids to a complete variant analysis.

## 2 Materials and Methods

### 2.1 Human Biological Samples

De-identified, remnant human biological FFPE from NSCLC patients were sourced from Discovery Life Sciences (Huntsville, AL), Dartmouth Hitchcock Medical Center (Lebanon, NH),and Cureline (Brisbane, CA). All samples enrolled in this study had no pathological selection criteria (Extended Table S7). FFPE samples were split into three groups based on “time in block” age (Table 1). Discovery Life Sciences and Dartmouth Hitchcock Medical Center isolated the nucleic acids (DNA and/or RNA) using validated in-house methods and performed initial quality control (QC) (quantification, sizing, and RNA quality assessment). The QC data, patient demographics, and clinical metadata for all samples are provided in Extended Table S7. Normal adjacent tissue (NAT) FFPE curls (Discovery) were combined in sets of three curls per tube and extracted with the AllPrep® DNA/RNA FFPE Extraction Kit (PN 80234, Qiagen, Germantown, MD). Isolated nucleic acids were quantified by Qubit4™ (Qubit dsDNA HS kit, ThermoFisher Scientific, Waltham, MA).

**Table 1.**
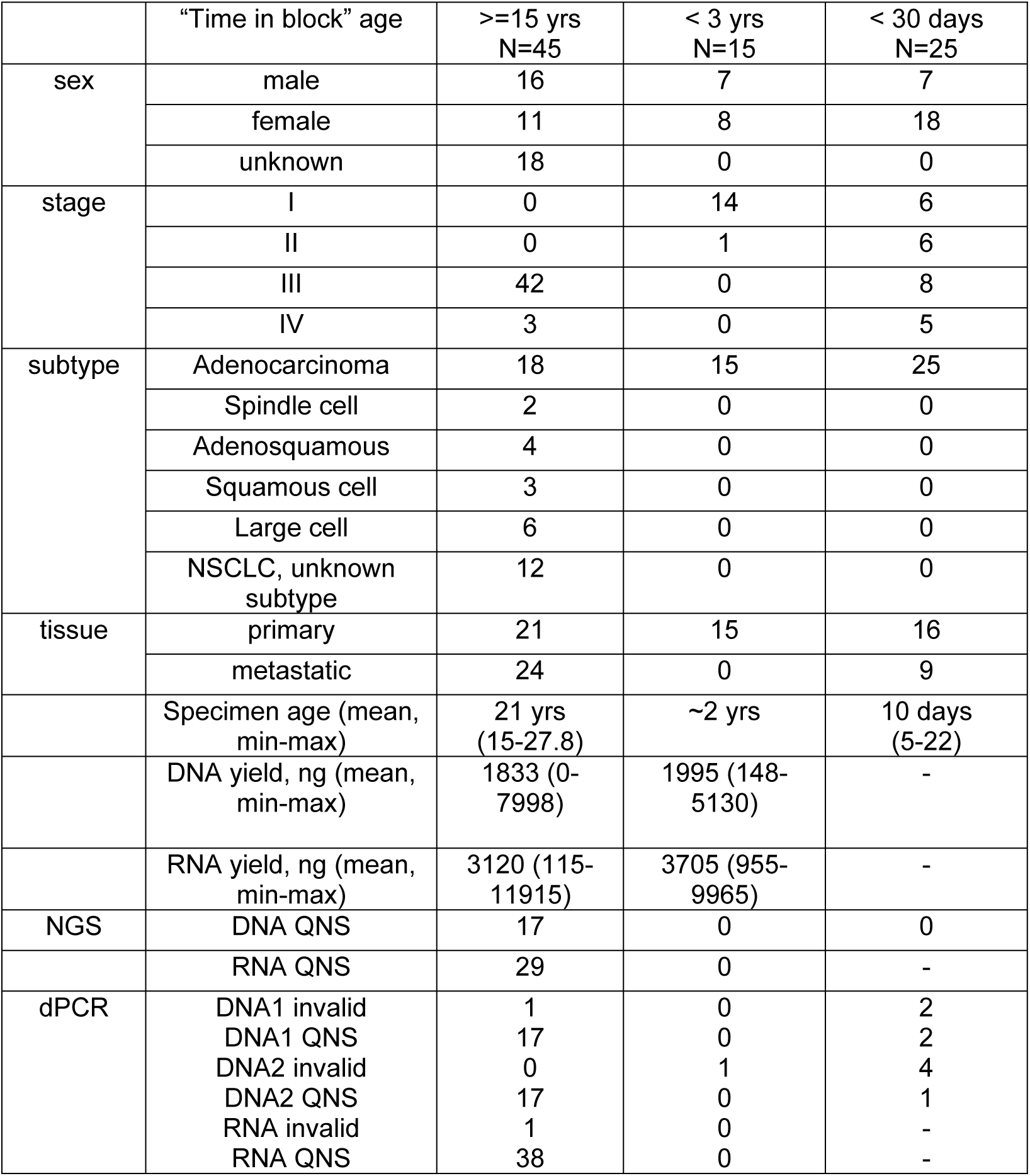
Human biological FFPE sample metadata and QC performance through sequencing and dPCR workflows.

|  | "Time in block" age | >=15 yrs<br>N=45 | < 3 yrs<br>N=15 | < 30 days<br>N=25 |
| --- | --- | --- | --- | --- |
| sex | male | 16 | 7 | 7 |
|  | female | 11 | 8 | 18 |
|  | unknown | 18 | 0 | 0 |
| stage | I | 0 | 14 | 6 |
|  | II | 0 | 1 | 6 |
|  | III | 42 | 0 | 8 |
|  | IV | 3 | 0 | 5 |
| subtype | Adenocarcinoma | 18 | 15 | 25 |
|  | Spindle cell | 2 | 0 | 0 |
|  | Adenosquamous | 4 | 0 | 0 |
|  | Squamous cell | 3 | 0 | 0 |
|  | Large cell | 6 | 0 | 0 |
|  | NSCLC, unknown subtype | 12 | 0 | 0 |
| tissue | primary | 21 | 15 | 16 |
|  | metastatic | 24 | 0 | 9 |
|  | Specimen age (mean, min-max) | 21 yrs<br>(15-27.8) | ~2 yrs | 10 days<br>(5-22) |
|  | DNA yield, ng (mean, min-max) | 1833 (0-7998) | 1995 (148-5130) | - |
|  | RNA yield, ng (mean, min-max) | 3120 (115-11915) | 3705 (955-9965) | - |
| NGS | DNA QNS | 17 | 0 | 0 |
|  | RNA QNS | 29 | 0 | - |
| dPCR | DNA1 invalid | 1 | 0 | 2 |
|  | DNA1 QNS | 17 | 0 | 2 |
|  | DNA2 invalid | 0 | 1 | 4 |
|  | DNA2 QNS | 17 | 0 | 1 |
|  | RNA invalid | 1 | 0 | - |
|  | RNA QNS | 38 | 0 | - |

### 2.2 Synthetic RNA via *in vitro* Transcription

The MEGAscript™ T7 Transcription Kit (PN AM1330, Life Technologies, Carlsbad, CA) was used according to the manufacturer’s protocol. First, custom DNA gBlocks with T7 promoter sequences (IDT, Coralville, Iowa) were created for each fusion variant (Table S2). The transcription reaction was set up with the following volumes using the MEGAscript T7 Transcription Kit: 2 µL ATP solution, 2 µL CTP solution, 2 µL GTP solution incubated, 2 µL UTP solution, 2 µL 10X Reaction Buffer, 8 µL IVT gBlock at 1E6 copies/µL, and 2 µL Enzyme Mix. The reaction mix was incubated at 37°C for 4 hours, and then 1 µL of TURBO™ DNase was added to the transcription reaction and incubated at 37°C for 15 minutes. *In vitro* transcription (IVT) products were initially evaluated for yield and purity by qPCR. This was done by creating two different reaction mixes, one with reverse transcriptase and one without, which tested for any remnant DNA before being used in contrived testing. Once the IVT fusion products were determined to not contain DNA, they were quantified with a singleplex dPCR assay for *ACTB*.

### 2.3 Amplitude-modulation dPCR assay construction

The primer-probe systems adopted one of three configurations: an allele-refractory mutation system (ARMS) with or without blocking oligonucleotides, a variant-sensitive probe, or an exon-specific design to identify exon-exon RNA fusion junctions (Figure 1). To begin, we synthesized and screened multiple primer-probe systems in singleplex using synthetic templates designed to represent a variant of interest. For the DNA-specific ARMS and variant-sensitive probe systems [20], the strandedness of the system (targeting Watson or Crick), the thermodynamics of the penultimate base pair mismatch, and the orientation with respect to nearby variant sites were considered during the design phase. Once systems were identified that worked well in singleplex and in pairwise duplex, the same principles of amplitude modulation in dPCR that have previously been demonstrated on qPCR [17] were applied. This approach allows multiple targets to be detected in the same color channel by tuning the reaction chemistry and probe concentrations, then applying Poisson statistics to interpret the observed dPCR data. Primer and probe concentrations were empirically optimized under multiple different concentration and thermal cycling conditions to achieve terminal fluorescent amplitude values that allowed for fluorescent intensity separation of all variants (Table S3). Due to the close genomic coordinate proximity of some of the DNA variants, the DNA targets were split into two separate wells to minimize cross-target amplification. For the RNA-specific fusion targets, a separate reaction included a reverse-transcription PCR step to generate cDNA. We also sought to incorporate knowledge of the prevalence and co-occurrence of certain biomarkers into the assay design. For example, to reduce the risk of calling errors that may be elevated in co-positive samples (e.g. *EGFR* L858R and *EGFR* Exon 19 deletion), prevalent variants were encoded in different color channels.

**Figure 1.**
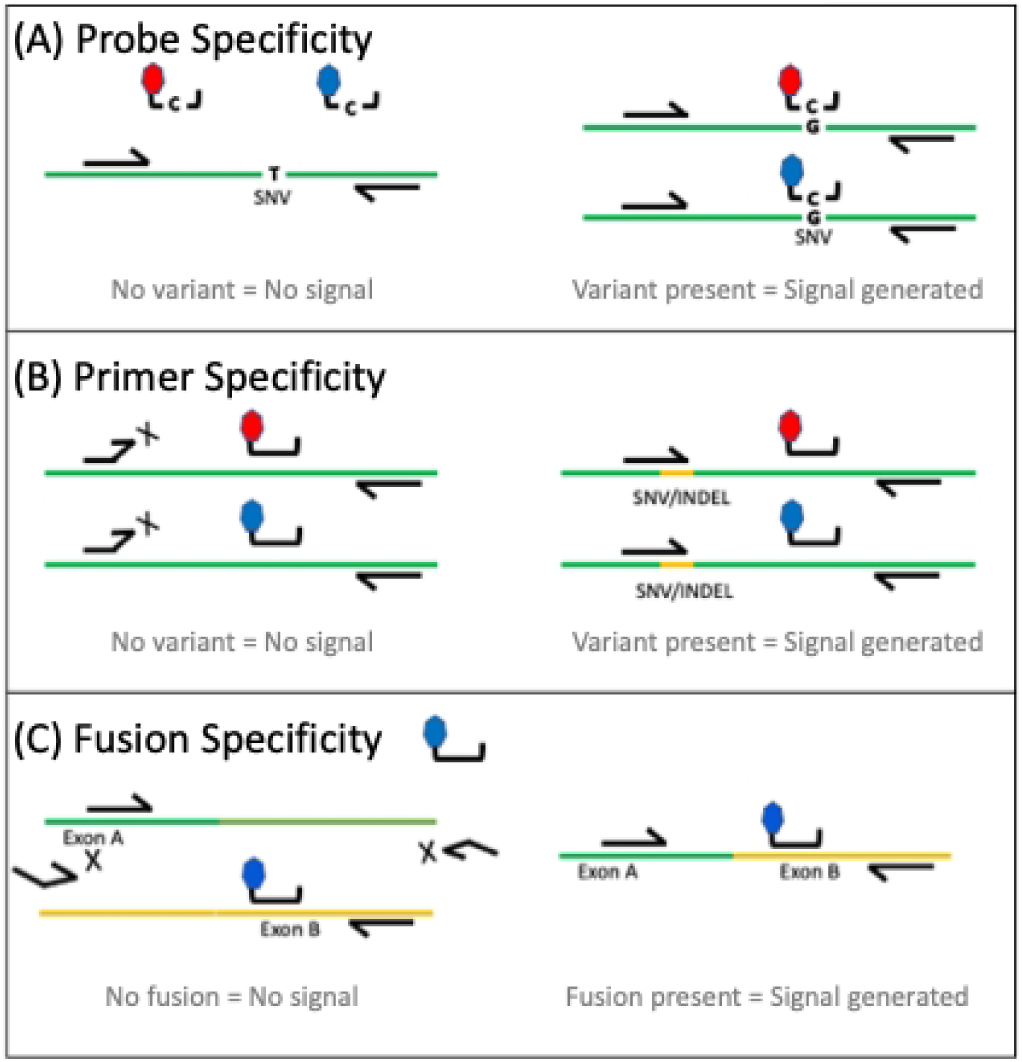
Three TaqMan primer/probe configurations are leveraged in the multiplex dPCR assay. A) One or two identical sequence probes, each with a different fluorophore / quencher pair (red and blue), hybridize specifically to the variant sequence and not to the wild type sequence. Probes are flanked by wild type locus-specific primers. B) ARMS primers specific for the SNV or indel of interest undergo 3’ extension if there is a perfect sequence match. One or two identical sequence probes complementary to wild type sequence can be labeled with different fluorophore / quencher pairs. C) RNA-based fusion assays designed against cDNA sequences whereby one primer targets one gene exon and a second primer and probe target the exon of the fusion partner gene.

Complete sets of multiplex primer-probe systems were prioritized based on four criteria: responsiveness to amplitude modulation, reaction efficiency (e.g. minimal dPCR “rain”), cross-reactivity due to proximity of targets, and specificity to discriminate between the variant and the wild-type sequences. The issue of “rain” refers to partitions with fluorescence amplitude that falls between the expected positive partition amplitude and the negative partition amplitude. For amplitude modulation PCR, the “rain” creates an additional issue where partitions belonging to a higher amplitude level (e.g. level 2 or 2i) are misclassified as a lower level (level 1 or 1i) thereby creating false positives in the lower-level windows and false negatives in the higher-level window. For some targets, locked nucleic acid (LNA) probe-based detection schemes [21, 22] had less interaction with wild-type DNA and produced less “rain”. Other primer and probe systems that had noticeably higher reaction efficiency (e.g. minimal “rain”) were assigned to higher intensity levels.

The nature of dPCR reduces the impact of nonspecific amplification events, as false positive signals are contained to a few partitions. However, it can still result in appreciable noise levels in the absence of target (Figure 2A, B). This led us to implement a multi-spectral encoding strategy for some targets to further improve performance (Figure 1A, B). Multi-spectral encoding relies on including two probes to the same target, each with a different fluorescent signature. This creates two independent probe hydrolysis events, thereby enhancing the signal above the noise created due to non-specific single probe hydrolysis. For example, the *EGFR* T790M system generated positive counts in the presence of wild-type genomic DNA (Figure 2A, B), and a similar number of counts in the presence of low copy number *EGFR* T790M variant (Figure 2D, E). However, when *EGFR* T790M is encoded in channel 5 as well as channel 1, the T790M positive counts are easily distinguished from the noise (Figure 2C, F). In another example, the channel 1 probe for *KRAS* G12C performed better than the channel 3 probe with extracted genomic DNA and by combining the two, a more distinct population of positive partitions are generated (Figure 3). Figure 4 shows the amplitude-modulation layout of the first multiplex DNA assay and example experimental data showing how amplitude modulation and multi-spectral encoding work together to resolve multiple variants in one well. Refer to Table S3 for a representative primer-probe formulation to achieve this assay layout, and Figure S1 for the terminal intensity layout for wells 2 and 3.

**Figure 2.**
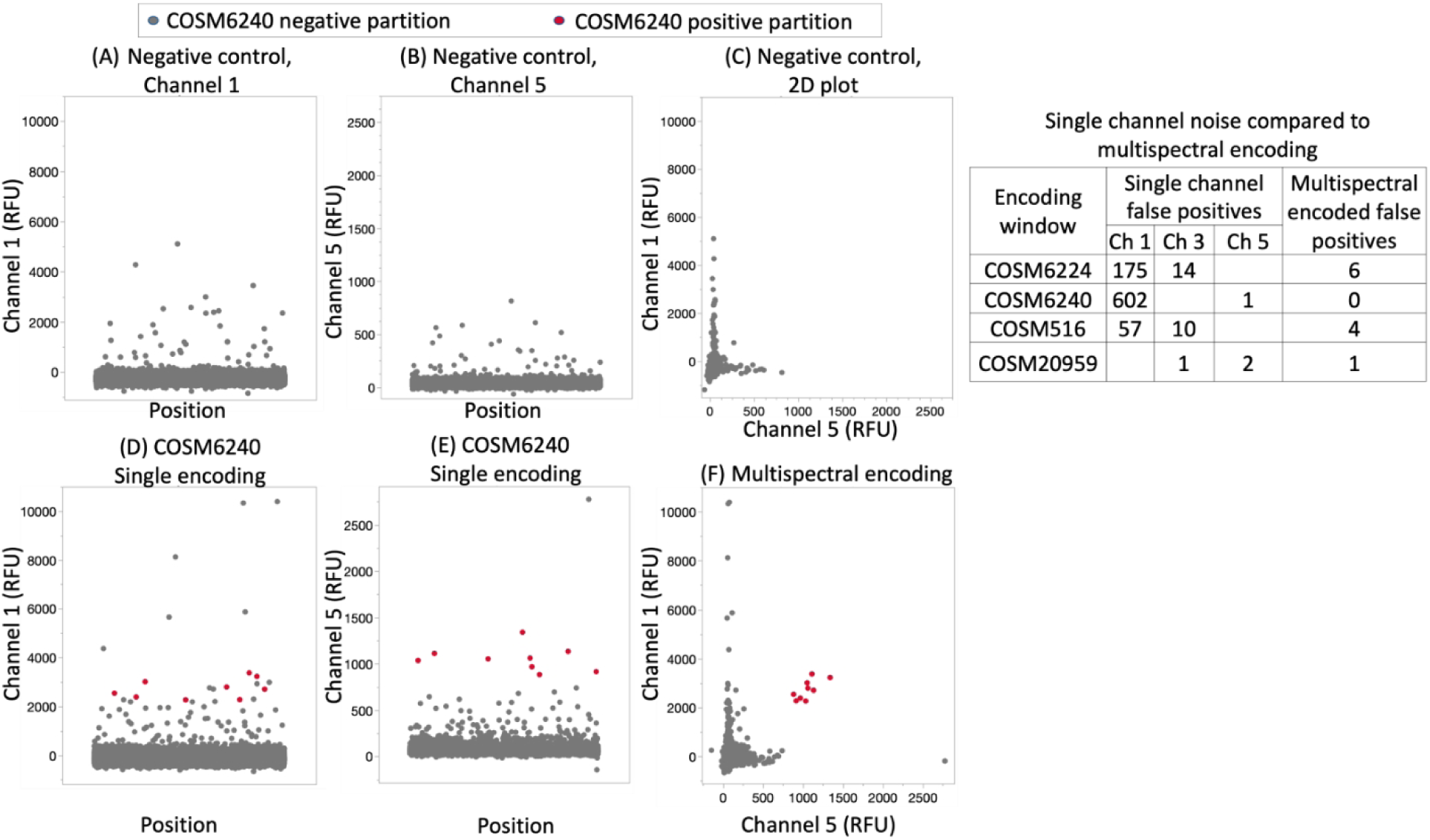
Multi-spectral encoding isolates background non-specific wild type amplification inherent to nucleic acid hybridization-reliant chemistry. Panels A), B) and C) show 1D and 2D plots in two channels for probe-based detection for COSM6240 (*EGFR* T790M). The primers and probes produce some non-specific amplification with background wild type DNA (N=6090 haploid genome copies). D, E) A contrived sample containing 0.25% COSM6240 synthetic copies in a background of wild-type DNA generates true positive signal in channel 1 that is indistinguishable from non-specific amplification. F) The same sample as in D) and E) leveraging multi-spectral encoding to isolate true positive partitions from non-specific amplification. The table on the right shows false positive counts arising within the call windows of each of four targets from four negative control samples.

**Figure 3.**
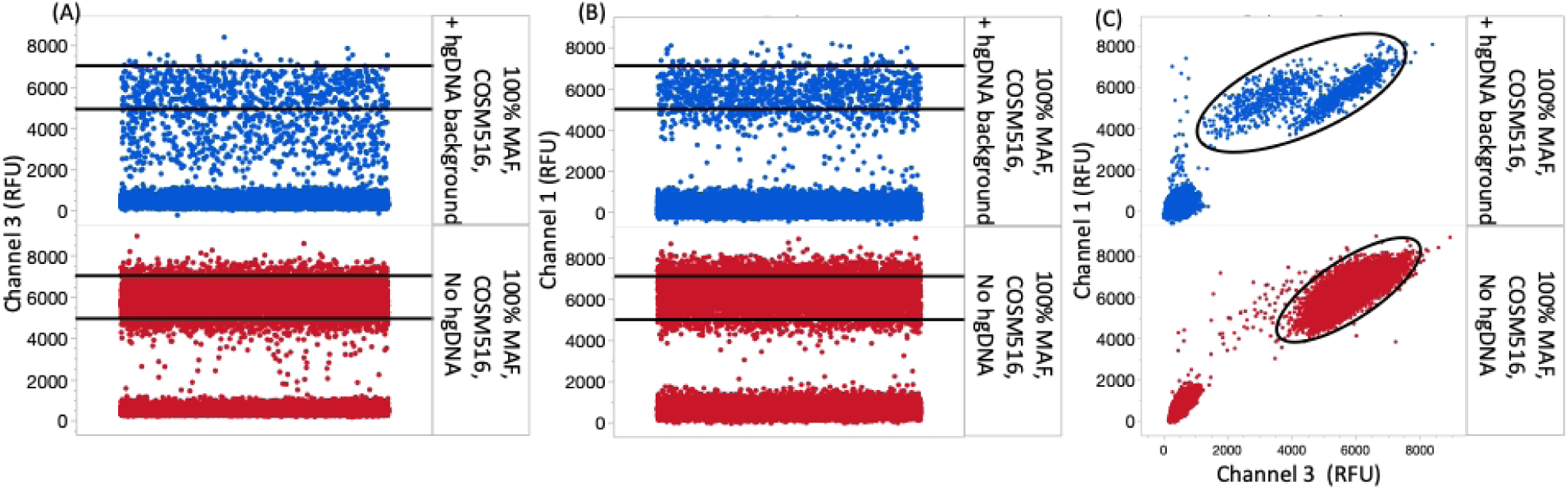
Multi-spectral encoding compensates for variable channel 3 and channel 1 probe performance. A, B) A channel 3 or 1 probe targeting COSM516 (*KRAS* G12C) in the presence of synthetic target and human genomic DNA (top) or synthetic target alone (bottom). (C) A mixture of channel 1 and channel 3-labeled COSM516 probes leads to a shift in the positive distribution away from the negative population in both the X and Y directions, reducing false positive partitions and consolidating true positives.

**Figure 4.**
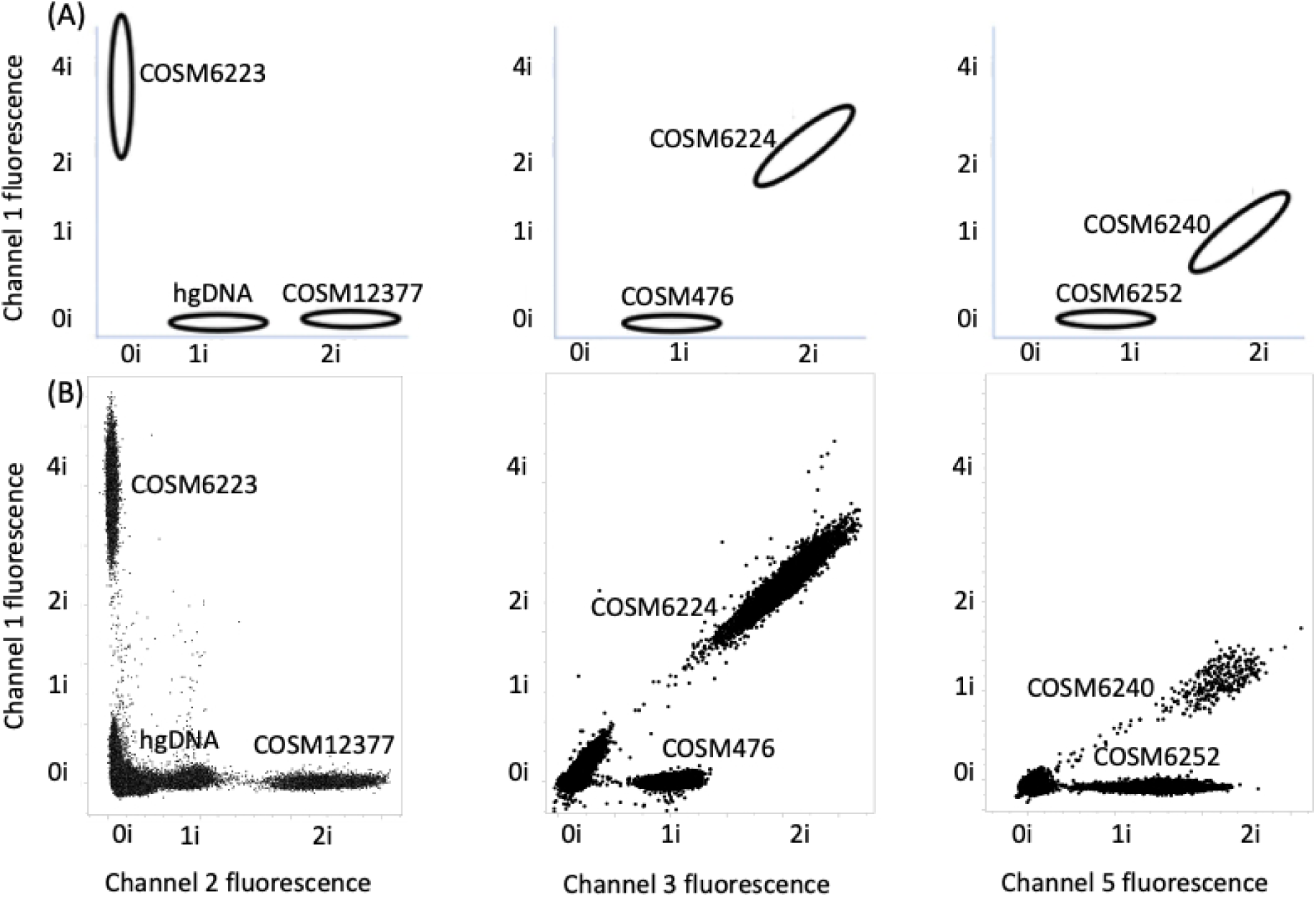
Amplitude modulation enables detection of multiple targets in a single channel. A) Approximate locations in two-channel space where each partition variant is expected for well #1 targets (Table 1). hgDNA refers to in-well positive control amplicon for *EGFR* Exon 2. B) Superimposed fluorescence scatterplots for synthetic targets profiled individually at 5000 copies for *EGFR* E746_A750del (COSM6223), *EGFR* Exon 20 H773dup (COSM12377), *EGFR* L858R (COSM6224), *EGFR* T790M (COSM6240), *EGFR* G719S (COSM6252), and *BRAF* V600E (COSM476). Negative controls are shown in Figure S2; a similar spectral layout for well #2 is shown in Figure S1.

### 2.4 Amplitude modulation digital PCR reaction setup and cycling (DNA)

DNA PCR reactions were set up using the following volumes: 2.4 µL 5X dPCR QuantStudio™ Absolute Q™ Master Mix (PN A52490, ThermoFisher Scientific, Waltham, MA), 2.9 µL oligonucleotide primer-probe mix (Table S3), and 6.7 µL of isolated genomic DNA. Contrived samples and natural specimen FFPE were tested at 4.18 ng/µl. Each dPCR reaction mix was vortexed three times for five second pulses, spun down in a microfuge, and 9 µL of the dPCR reaction mix was added to each well of a QuantStudio Absolute Q MAP16 Plate Kit (PN A52865, ThermoFisher Scientific, Waltham, MA). Next, 12 µL of QuantStudio Absolute Q Isolation Buffer (PN A52730, ThermoFisher Scientific) was added to each well on top of each reaction mix. The final quantity of genomic DNA that makes it into the system, as part the 9 µL input, is 21 ng for contrived and FFPE samples. The wells were sealed with QuantStudio Absolute Q strip caps (PN 332101, ThermoFisher Scientific). All testing was conducted on one of two QuantStudio Absolute Q Digital PCR Systems (Thermo Fisher Scientific). Thermal cycling was performed as follows: (1) Preheating at 96°C for ten minutes, (2) 35 cycles consisting of denaturing (96°C, 15 seconds), followed by annealing/extension (58°C for 30 seconds). Terminal fluorescence intensity data was collected in all four available color channels. Along with the reaction mixes, every plate included a positive control (gBlocks of synthetic targets in each color channel) and a negative control (consisting of only human genomic DNA background). Positive control primers for *EGFR* Exon 2 (DNA) were included in each well, respectively. Primer and probe sequences are described in Table S1 (Integrated DNA Technologies, Inc. (Coralville, IA) and ThermoFisher Scientific (Waltham, MA)).

### 2.5 Amplitude-modulation digital PCR reaction setup and cycling (RNA)

RNA dPCR reactions were set up using the following volumes: 2.4 µL 5X dPCR QuantStudio Absolute Q Master Mix (PN A52490, ThermoFisher Scientific, Waltham, MA), 2.4 µL 5X primer-probe mix (Table S3), 0.6 µL reverse transcriptase (PN M0368S, New England Biolabs, Ipswich, MA), 5 µL RNA sample (1-3 ng total RNA), and 1.6 µL 1X TE Buffer (pH 8.0, Low EDTA (Tris-EDTA; 10 mM Tris base, 0.1 mM EDTA)) (PN 786-150, G-Biosciences, St. Louis, MO). Each dPCR reaction mix was then vortexed three times for five second pulses, spun down in a microfuge, and 9 µL of the dPCR reaction mix was added to each well of a QuantStudio Absolute Q MAP16 Plate (PN A52865, ThermoFisher Scientific). Next, 12 µL of QuantStudio Absolute Q Isolation Buffer (PN A52730, Thermo Fisher Scientific) was added to each well on top of the reaction mix. The wells were sealed with QuantStudio Absolute Q strip caps (PN 332101, ThermoFisher Scientific). All testing was conducted on QuantStudio Absolute Q Digital PCR Systems (ThermoFisher Scientific). Thermal cycling was performed as follows: (1) Reverse Transcription at 50° C for 15 minutes, (2) Preheating at 95°C for 10 minutes, (3) 40 cycles consisting of denaturing (95°C for 10 seconds), followed by annealing/extension (58°C for 1 minute). Terminal fluorescence intensity data was collected for all 4 available color channels. Along with the reaction mixes, every plate included a positive control (gBlocks of synthetic targets in each color channel) and a negative control (consisting of only isolated FFPE total RNA background). Positive control primers for *ACTB* (RNA) were included in each well, respectively. Primer and probe sequences are described in Table S1 (Integrated DNA Technologies, Inc. (Coralville, IA) and Thermo-Fisher Scientific (Waltham, MA)).

### 2.6 Contrived DNA and RNA sample assembly

Contrived FFPE samples were created by combining synthetic DNA gBlocks (average size = 400 nt, containing either reference sequence or variant of interest, from IDT, Coralville, Iowa) with 21 ng of extracted healthy (negative) human FFPE DNA at six different variant fractions ranging from 60-2300 copies (1-40% VAF). The contrived FFPE RNA samples were created by combining the fusion IVT RNAs with the negative extracted FFPE RNA at a range of copy numbers: 5000, 7500, 10000, 11250 while the negative extracted FFPE RNA remained constant at 5000 copies (Table S4).

### 2.7 Variant calling from amplitude modulated digital PCR data

Once the oligo sequences and concentrations were set for each assay, a run was conducted with each genomic target present in singleplex in two replicate wells. For each target, positive partitions were identified using an amplitude cutoff which was established by testing each target in the assay individually, and the mean and covariance of positive partition amplitudes were calculated across all four channels. The mean and covariance of partition amplitudes for all possible target combinations were predicted by assuming amplitudes would add linearly. This set of analyses generated “expected” target amplitudes, which were used to classify partitions across all other experiments. These singleplex runs were also used to characterize the crosstalk levels of each dPCR instrument, and this crosstalk was subtracted out in all multiplex runs.

Each sample plate run contained at least one negative control well, which only had the internal *EGFR* Exon 2 control target present, and at least one positive control well, which had multiple synthetic targets present that would generate signal in each channel. These controls were used to perform three plate-wide corrections. First, the negative control well was used to determine the mean amplitude of partitions positive for the internal control; if this was different from the expected location, then the expectation for that target was scaled for the rest of the plate. Similarly, the positive control well was used to determine the mean amplitude of partitions which were positive in each individual channel. If a given channel differed from its expected level, the ratio between observed and expected mean was used to scale the expected amplitude for all targets in that channel. Finally, the negative control well was re-analyzed to determine how many partitions were positive for targets other than the internal control target. These levels were used to determine an expected level of spurious amplification which occurs in the absence of target material. This set of corrections was performed on a plate-by-plate basis to correct for any differences from run to run.

After these plate-wide corrections, non-control wells were analyzed to determine target counts. Partition classification was performed using the Mahalanobis distance metric: for a partition with the 4-dimensional amplitude vector 𝑥, its Mahalanobis distance to a target with expected mean amplitude 𝜇 and covariance matrix 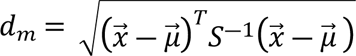. This is effectively the same as classic Euclidian distance but scaled by the covariance of the expected target amplitude; this corrects for the fact that some targets generate point clouds with inherently wider spread than others. Each partition is assigned to the target or target combination to which it has the lowest Mahalanobis distance. Analyzing all partitions in this manner results in a count of positive partitions for each target, which is converted into a target concentration using Poisson statistics. The expected level of spurious amplification was then subtracted to yield a final concentration for each target.

For the contrived and human biological sample experiments, DNA samples with *EGFR* Exon 2 copy numbers below 1000 copies per reaction were empirically determined to be Quantity Not Sufficient (QNS) and excluded from the performance calculations. Similarly, RNA samples with *ACTB* copy numbers below 1000 copies per reaction were determined to be QNS. To quantitatively determine which samples exhibited abnormal results, wells were labeled invalid and excluded from the analysis if they had a coefficient of variation in the reference channel across all partitions of greater than 15% (Figure S3b). Additionally, if a well had greater than 100 partitions with signals less than 6000 relative fluorescent units in the reference channel, it was determined to be invalid and excluded from the analysis. These exclusions led to an observed per-well failure rate of ∼4.95% (33/666 total reactions) on both instruments. One of the main failure modes was images with dark patches in the QC array (Figure S3a), which could be due to optical or flow issues in the instrument. The performance of the chemistry and algorithm was determined on the contrived DNA samples down to 1% VAF and the contrived RNA samples down to 5000 total fusion copies (Table 1 and Table S4). Receiver operating characteristic (ROC) analysis was performed on the complete DNA and RNA contrived data sets to identify the optimal threshold for each target to separate positive and negative contrived samples. The ROC analysis used the ratio of the target to the in-well positive control (*EGFR* Exon 2 and *ACTB* for the DNA and RNA assays, respectively) as the predictor. The calculations were performed using the R software package pROC [23, 24]. These optimized thresholds were used to calculate the performance of the clinical sample data sets.

### 2.8 Parallel comparator testing

DNA and RNA isolated from the Discovery and Cureline FFPE clinical samples were parallel processed through Discovery Life Sciences’ QiaSeq MultiModal panel (64 DNA genes and 6 primary genes for RNA fusions, recommended input mass of 200 ng DNA and 200 ng RNA with at least DV20%). Data were processed through Qiagen’s CLC Workbench bioinformatics workflow to generate variant call files and reports. DNA isolated from the Dartmouth Hitchcock samples were processed using Ion AmpliSeq™ Cancer Hotspot Panel v2 and TruSight Tumor 170. Data processing was performed using the Torrent Suite and the TruSight Tumor 170 v1.0 Local App respectively. Sequencing summary statistics are provided in Extended Table S8. Samples were considered indeterminate and excluded from the clinical concordance analysis if the sample did not generate at least 20 reads for a particular target region.

To improve comparator confidence in 31 RNA samples with low read counts for *ALK* and/or *RET* transcripts (<100 RPKM), we sought to run an additional fusion comparator that was commercially available using digital droplet PCR. RNA from the FFPE clinical samples were processed through BioRad mRNA ddPCR fusion assays for *RET* (ID dHsaEXD81378442, Bio-Rad, Hercules, CA), *ROS1* (ID dHsaEXD73338942, Bio-Rad, Hercules, CA), and *ALK* (ID dHsaEXD86850342, BioRad, Hercules, CA). First, the clinical FFPE RNA samples were converted to cDNA using the BioRad cDNA synthesis kit according to the manufacturer’s instructions (PN 1725037, Hercules, CA). cDNA was then quantified by Qubit4 (ThermoFisher Scientific, Waltham, MA). Three ddPCR reactions were set up for each of the mRNA fusion assays with the following volumes: 10 µL 2x ddPCR Supermix for Probes (no dUTP) (PN 186-3023, Bio-Rad, Hercules, CA), 1 µL 20x mRNA Fusions Assay, 1 µL 20x *GUSB* Reference Assay (ID dHsaCPE5050189, Bio-Rad, Hercules, CA), 6 µL cDNA, and 4 µL nuclease-free water (PN 10977015, Invitrogen, Waltham, MA). Each ddPCR reaction mix was added to a 96-well PCR plate (PN 12001925, Bio-Rad, Hercules, CA), sealed with a PX1 PCR Plate Sealer (Bio-Rad, Hercules, CA), and then vortexed 3 times for 10 second pulses, and spun down in a microfuge. The ddPCR fusion reactions were first run on the Bio-Rad Automated Droplet Generator (Bio-Rad, Hercules, CA) according to the manufacturer’s instructions. Once the droplets were generated, a new reaction plate was generated and sealed with a PX1 PCR Plate Sealer. This new reaction plate was transferred to a C1000 Touch Thermal Cycler (Bio-Rad, Hercules, CA) and run at the following conditions: (1) Preheating at 95°C for 10 minutes, (2) 40 cycles of denaturing (94°C for 30 seconds), followed by annealing/extension (55°C for 1 minute), and 3) enzyme deactivation at 98°C for 10 minutes. Lastly, the reaction plate was run on the QX200 Droplet Reader (Bio-Rad, Hercules, CA) according to manufacturer’s instructions. The results were analyzed by *GUSB* counts to determine valid/invalid samples. The Bio-Rad fusion assays were then benchmarked to our fusion assay by running each of them with a titration of IVT products (0, 10, 50, 100, 500 copies) in a 1 ng background of RNA cell line reference (PN 4307281 Applied Biosystems, Waltham, MA).

## 3 Results

### 3.1 Contrived sample and commercial reference performance

After removing invalid samples (n=40 DNA, n=7 RNA), a total of 293 FFPE DNA and 314 FFPE RNA contrived reactions, each containing one or more variants at a range of variant allele frequencies (Table S4), were characterized on the multiplexed assay. These samples were constructed with no *a priori* knowledge on the assay performance, as we sought to understand calling accuracy at both high and low VAFs using a custom algorithm designed to automatically classify each digital partition (see Methods). With the parameters optimized for the contrived sample set, the algorithm calling gave results in agreement with the contrived sample composition: for the contrived FFPE DNA and RNA targets, a 94% PPA / 99% NPA and 100% PPA / 97.9% NPA, respectively (Table 2 and Table S6).

**Table 2.** Contrived human biological sample performance. Algorithm performance on all contrived samples at >=1% VAF. Algorithm parameters were optimized on this same sample set as described in the Methods.

| Target | True Positive | False Negative | False Positives | True Negative | PPA | NPA |
| --- | --- | --- | --- | --- | --- | --- |
| *EGFR* L858R | 79 | 0 | 0 | 124 | 100% | 100% |
| *EGFR* T790M | 52 | 8 | 7 | 136 | 86.7% | 95.1% |
| *EGFR* E746_A750del | 48 | 0 | 1 | 154 | 100% | 99.4% |
| *BRAF* V600E | 6 | 0 | 0 | 197 | 100% | 100% |
| *EGFR* G719S | 40 | 12 | 0 | 151 | 76.9% | 100% |
| *EGFR* H773dup | 17 | 0 | 0 | 186 | 100% | 100% |
| *KRAS* G12C | 20 | 0 | 1 | 69 | 100% | 98.6% |
| *ERBB2* Y772_A775dup | 15 | 0 | 0 | 75 | 100% | 100% |
| *EGFR* S768I | 9 | 0 | 0 | 81 | 100% | 100% |
| *EGFR* L861Q | 9 | 0 | 0 | 81 | 100% | 100% |
| Total | 295 | 20 | 9 | 1254 | 93.7% | 99.3% |
| *EML4-ALK* | 62 | 0 | 3 | 249 | 100% | 98.8% |
| *MET* Exon14 | 47 | 0 | 0 | 267 | 100% | 100% |
| *KIF5B-RET* | 73 | 0 | 0 | 241 | 100% | 100% |
| *CD74-ROS1* | 76 | 0 | 0 | 238 | 100% | 100% |
| *TMP3-NTRK1* | 56 | 0 | 24 | 234 | 100% | 90.7% |
| Total | 314 | 0 | 27 | 1229 | 100% | 97.9% |

The assay generated a total of 20/1578 (1.3%) false negative calls and 9/1578 (0.6%) false positive calls on the contrived DNA samples, which may be partly driven by chemistry and partly by instrument noise. For example, the majority of these false negative DNA calls were associated with the *EGFR* G719X target. This primer/probe system was one of the noisiest, likely because it targeted three variants with three different variant-specific primers at the same codon and required a blocker to suppress the wild-type signal. The *EGFR* G719X assay was not multi-spectrally encoded, which could have significantly reduced the non-specific calls and allowed for higher amplitudes to increase sensitivity.

We further sought to assess the analytical accuracy of the RNA assay using an external reference standard (SeraCare). Here, three of the fusion reportables (*ALK*, *ROS1*, and *MET* Exon 14 skipping) were tested with the multiplex dPCR assay and found to generate copy estimates in strong agreement with the Certificate of Analysis concentration (Table S5). An additional comparison of the multiplex dPCR assay against three commercially available singleplex fusion assays for *ALK*, *RET*, *ROS1* also demonstrated similar levels of performance, with a sensitivity to detect 100 or fewer IVT RNA molecules (Figure S4).

### 3.2 Human biological sample performance

Consistent with prior reports on the impact of FFPE storage time on DNA fragment length [25], 17/45 FFPE samples that were >15 years old did not yield DNA of sufficient quality to generate libraries for sequencing or dPCR analysis. All the 40 FFPE samples that were <3 years old yielded sufficient DNA and RNA for sequencing (>200 ng of DNA and RNA). However, prioritizing material for sequencing left three samples with insufficient material for subsequent dPCR testing. After filtering for samples with both passing dPCR calls and sufficient NGS read data at each target position, the assay achieved a 100% PPA and 98.5% NPA on the human biological FFPE DNA samples (n=38), and a 100% NPA on the FFPE RNA samples (n=31) (Table 3 and Table S6a).

**Table 3.** Nucleic acids from human biological NSCLC samples were isolated and underwent QC as described in the Methods section. Results are shown for samples passing both NGS and dPCR QC and count criteria.

| Target | True Positive | False Negative | False Positives | True Negative | PPA | NPA |
| --- | --- | --- | --- | --- | --- | --- |
| *EGFR* L858R | 9 | 0 | 2 | 27 | 100% | 93.1% |
| *EGFR* T790M | 1 | 0 | 0 | 37 | 100% | 100% |
| *EGFR* E746_A750del | 10 | 0 | 1 | 22 | 100% | 95.7% |
| *BRAF* V600E | 0 | 0 | 0 | 35 | NA | 100% |
| *EGFR* G719S | 0 | 0 | 2 | 36 | NA | 94.7% |
| *EGFR* H773dup | 0 | 0 | 0 | 36 | NA | 100% |
| *KRAS* G12C | 0 | 0 | 0 | 32 | NA | 100% |
| *ERBB2* Y772_A775dup | 0 | 0 | 0 | 32 | NA | 100% |
| *EGFR* S768I | 0 | 0 | 0 | 34 | NA | 100% |
| *EGFR* L861Q | 0 | 0 | 0 | 37 | NA | 100% |
| Total | 20 | 0 | 5 | 329 | 100% | 98.5% |
| *EML4-ALK* | 0 | 0 | 0 | 20 | NA | 100% |
| *MET* Exon14 | 0 | 0 | 0 | 20 | NA | 100% |
| *KIF5B-RET* | 0 | 0 | 0 | 20 | NA | 100% |
| *CD74-ROS1* | 0 | 0 | 0 | 20 | NA | 100% |
| *TMP3-NTRK1* | 0 | 0 | 0 | 20 | NA | 100% |
| Total | 0 | 0 | 0 | 100 | NA | 100% |

Of the 28 DNA and 16 RNA samples >15 years old that generated sequencing data, we observed highly variable sequencing coverage across the variant loci interrogated by the dPCR assay (Extended Table S8). This appears to have contributed to five samples with clinical annotation of *EGFR* Exon 19 del+ (based on prior sequencing or PCR assays on sister blocks) where the dPCR assay detected *EGFR* E746_A750del (COSM6223), and NGS re-sequencing failed to detect a variant due to lack of coverage in Exon 19. Similarly, four samples were detected to be positive for *KRAS* G12C by dPCR, and three had associated clinical annotation of *KRAS*+, but they failed to generate sequencing data due to insufficient quantity of nucleic acid for library preparation (Table S6b). One sample was detected to be positive for *EGFR* H773dup but gave zero aligned reads in *EGFR* Exon 20. For the 40 DNA samples < 3 years old, one sample (DH-EGFR-048) was called dPCR positive for *EGFR* G719X that was not detected by NGS. Here, the comparator sequencing assay was validated for detection down to 5% G719X variant frequency, while the amplitude-modulation dPCR assay measured it at 2.0% VAF, suggesting it may have been missed by sequencing. Unfortunately, discordant resolution could not be performed on these samples as additional nucleic acid could not be obtained. Taken together, these results highlight the potential value of a dPCR assay that is compatible with lower input mass and yet still has high sensitivity to generate actionable information from degraded or low yielding samples.

### 3.3 Multi-spectral encoding improves TaqMan assay specificity

Based on the performance of single-probe TaqMan systems, we implemented multi-spectral encoding for *EGFR* L858R, *EGFR* T790M, *ERBB2* Y772_A775dup, and *KRAS* G12C (Figure 4 and S1). In the absence of multi-spectral encoding, the average single-channel background noise for these four targets was 108 positive partitions, as measured by running wild-type genomic DNA (Figure 2). With multi-spectral encoding, however, the average background noise for these four targets was reduced to an average of 3 positive partitions. Multi-spectral encoding thus allowed for the accurate counting of these targets down to as few as 9 molecules (Figure 2).

## 4 Discussion

There are a growing number of targets and associated molecular testing methodologies to interrogate NSCLC molecular tumor profiles, ranging from single gene qPCR tests [26], easy to use cartridge-based systems [27], to comprehensive genomic profiling assays [28]. Here, we describe a first-of-its-kind, proof-of-concept assay that combines the speed and simplicity of a PCR test with the breadth of actionable coverage and sensitivity of a multi-gene sequencing-based test. One of the key challenges with developing a highly multiplexed oncology-focused PCR assay is being able to separately and specifically report variants that are in very close physical proximity (e.g. *EGFR* L858R and *EGFR* L861Q, only separated by two codons). Primer and probe systems for one variant can inadvertently interact with the primer and probe systems for the other, leading to false positive signal generation. Here, we mitigated these interactions by either separating out proximal variants into separate wells, or by leveraging target-specific probes and a common, wild type amplicon that spans multiple targets. Additionally, we incorporated multi-spectral signal encoding to suppress wild type amplification noise that becomes increasingly more challenging in high multiplex PCR mixtures.

For a subset of the >15 year old DNA FFPE cases, there was insufficient nucleic acid available to proceed with library preparation and sequencing (N=17/45, 38%, Table 1) , or there was insufficient amplicon coverage across all actionable genomic positions to enable confident calls for all reportables (N=10/45, 22%). Amplicon coverage is a known issue for targeted sequencing panels and can be driven by a combination of isolation methods, hybridization capture probe locations, DNA fragment lengths, DNA input amount, and sequencing alignment workflows [29]. Here the issue was particularly acute, given the age of a large fraction of the samples. The multiplex dPCR assay, less constrained by DNA quality and input mass requirements for sequencing, was able to generate a valid result for 22 DNA samples that had insufficient DNA for sequencing or had coverage gaps (Table S6b). This highlights an important potential use case for a multiplex dPCR panel: for samples that are intended to be sequenced but have insufficient material, reflexing to a dPCR assay may be able to yield actionable information without the need for additional biopsy. To support this hypothesis, future work will explore the dPCR assay performance on additional FFPE sample types, including needle core biopsies and fine needle aspirates, where input mass is particularly challenging.

While some sequencing-based assays detect fusions through DNA measurements by attempting to identify specific breakpoints within introns, this can be computationally challenging and highly dependent on sequencing coverage [30]. For this reason, we selected a sequencing comparator that leverages RNA-seq, which like our assay, makes calls by detecting the presence of fusion exon-exon junctions. However, despite having a ∼50 ng total RNA input, we noticed that three of the RNA gene targets (*MET*, *NTRK1*, and *ACTB*) had low wild type expression levels (<100 RPKM) across all samples tested, which suggests some combination of pre-analytic and/or biological factors can create greater challenges for RNA-based fusion variant detection. The low read count held true for both the older (>15 year) and younger (<3 year) FFPE samples (Extended Table S8). To investigate whether the low counts were specific to sequencing, we evaluated the human biological samples with a second fusion comparator: three commercially available ddPCR singleplex fusion assays for *ALK*, *RET*, and *ROS1* (BioRad). We first verified the performance of the ddPCR BioRad assays by titrating the previously generated IVT products, and then proceeded with re-testing the human biological RNA samples. Of the N=60 RNA samples tested across the three ddPCR assays (1 ng total RNA for each assay), N=75/180 (42%) assays failed on the BioRad ddPCR assay due to low reference gene *GUSB* counts. In contrast, the amplitude modulation dPCR assay had only 6/60 (10%) assay failures due to reference gene copies with the same approximate (1.5-3 ng) of total RNA input. This highlights the importance of selecting suitable reference controls given pre-analytic and biological factors, as well as assay input mass.

In summary, the performance of the dPCR assay was evaluated using a mix of contrived and human biological NSCLC samples to assess performance. The contrived samples allowed testing across all variants and reportables at a range of VAFs, and enabled algorithm development and optimization. The assay also successfully detected many of the common DNA variants in NSCLC human biological samples, including variants present in samples that were not sufficient for NGS. While this assay nor the comparator assays did not detect any rare DNA variants or any RNA fusion positive samples, this is not surprising given the sample size and the low prevalence of rare variant and fusions (1-4% of NSCLC patients) [31, 32, 33, 34]. To further establish the potential of amplitude modulation digital PCR in NSCLC testing, additional work is needed to 1) expand the inclusivity of the assay for insertion, deletion, and fusion variants, 2) better understand the relationship between sample input, quality and performance, and 3) test the methods on a larger sample set containing representative rare variants and fusion positive samples.

## 5 Conclusions

Amplitude modulation and multi-spectral encoding enables laboratories to increase the amount of information and decrease noise in digital PCR reactions. Here, we illustrate how a 27-variant tumor profiling assay can be constructed for actionable biomarkers with a performance commensurate to next generation sequencing, with the benefit of compatibility with lower input mass samples. These chemical and computational approaches may help enable low-cost, fast turnaround, accessible assays in the future.

## Author Contributions

BL, KM, HS, JA, MW, AR, JS designed experiments; BL, KM, HS, JA, JS performed experiments and analyzed data, DY and LJ wrote the algorithms and analyzed data, and BL, KM, HS, LJ, DY, JS wrote the manuscript, with JA, DG, GT, AR input.

## Abbreviations

dPCR: digital polymerase chain reaction
FFPE: formalin-fixed paraffin embedded
HDPCR™: high definition polymerase chain reaction
Indel: insertion / deletion
IVT: in vitro transcription
LNA: locked nucleic acid
NAT: normal adjacent tissue
NGS: next generation sequencing
NPA: negative percent agreement
NSCLC: non-small cell lung cancer
PPA: positive percent agreement
RPKM: reads per kilobase of transcript per million reads mapped
SNV: single nucleotide variant
VAF: variant allele frequency
QNS: quantity not sufficient

## Supporting information

Table S8 Molecular Data

Table S7 Molecular Data

## Supplementary Material

**Figure S1.**
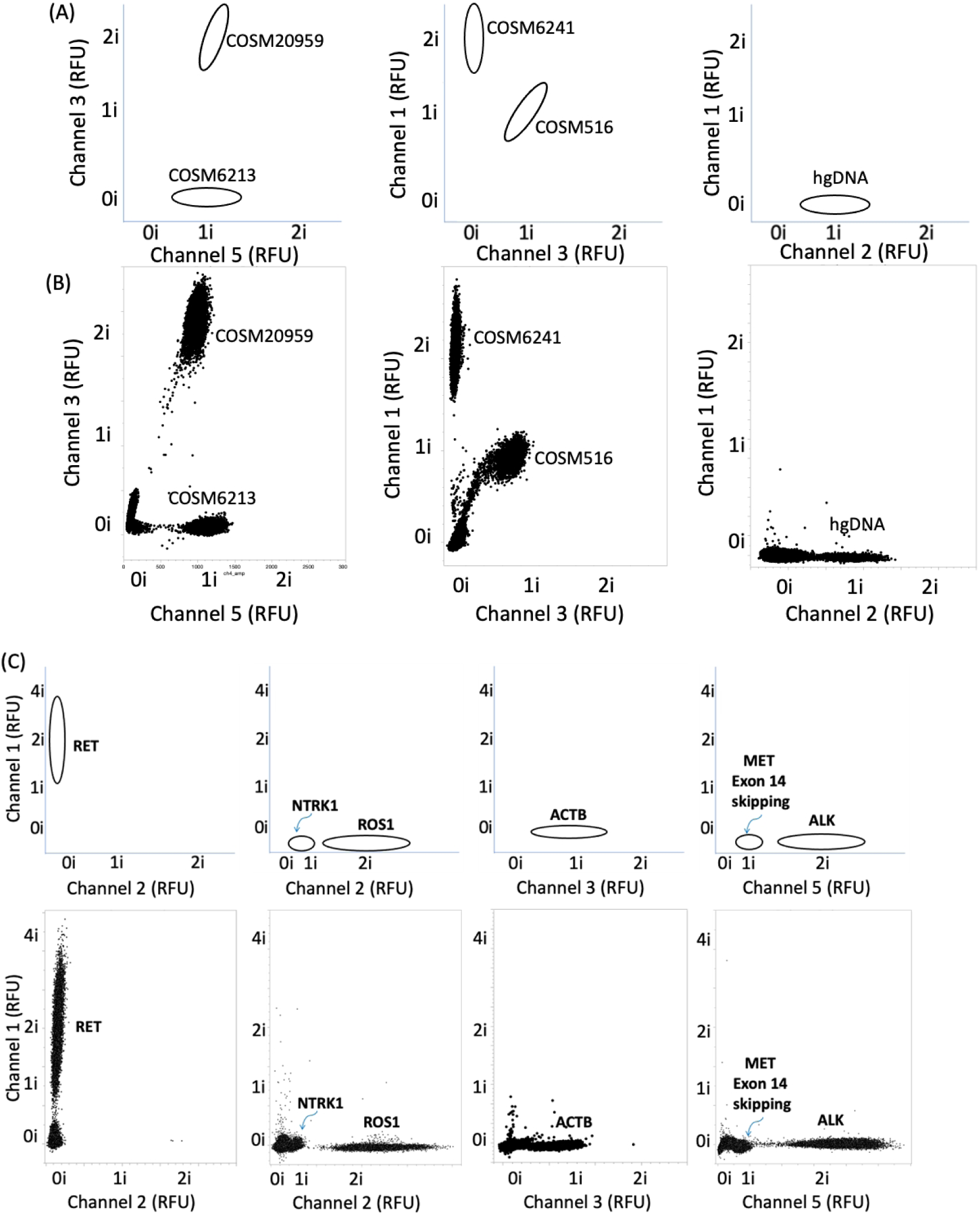
A) Illustrative locations on two-channel plot space where each variant-positive partitions are assigned for well #2 targets. B) Superimposed fluorescence scatterplots for synthetic targets profiled individually for *EGFR* S768I (COSM6241), *ERBB2* (COSM20959), *EGFR* L861Q (COSM6213), and *KRAS* G12C (COSM516). C) Approximate gate locations for RNA well #3 and example scatterplot showing the targets landing in the associated call window. RFU = Relative Fluorescence Units.

**Figure S2.**
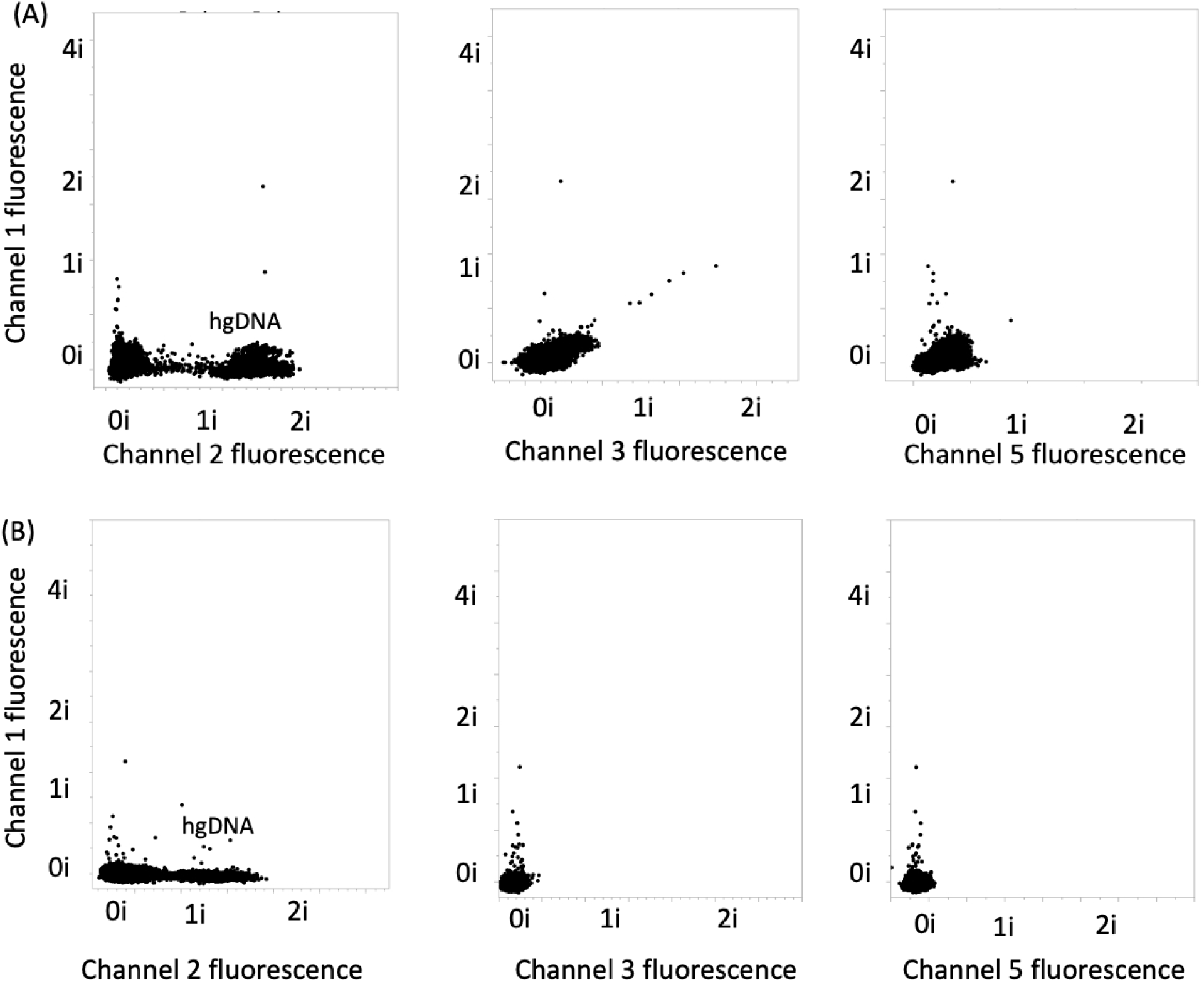
(A) Negative control (only human genomic DNA present) data for the plots shown in Figure 4. (B) The negative control for well #2.

**Figure S3a:**
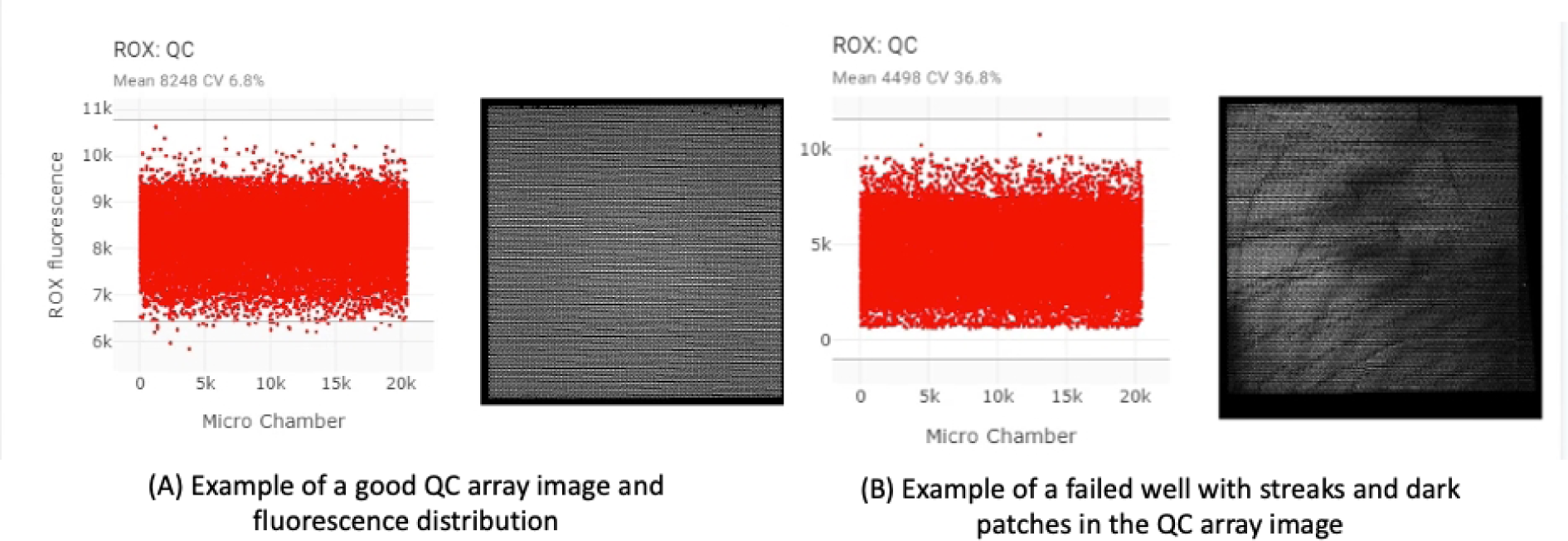
Comparison of the QC data and raw image of a valid Absolute Q well (left) and a failed well (right).

**Figure S3b:**
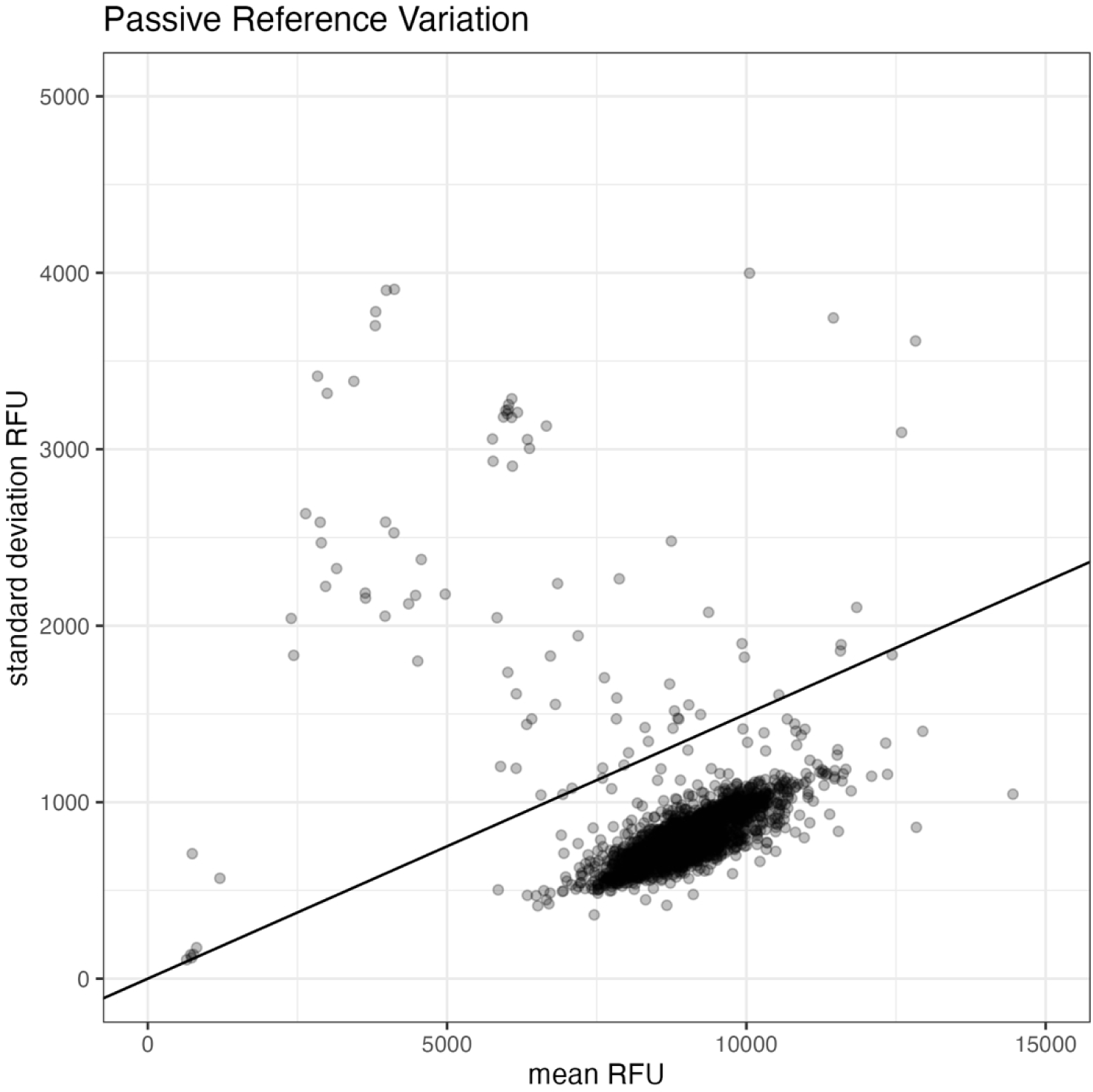
Variation of the passive reference signal for every well in both the clinical and contrived data sets. The line represents the 15% coefficient of variation threshold; all wells above this threshold were excluded from the analysis.

**Figure S4:**
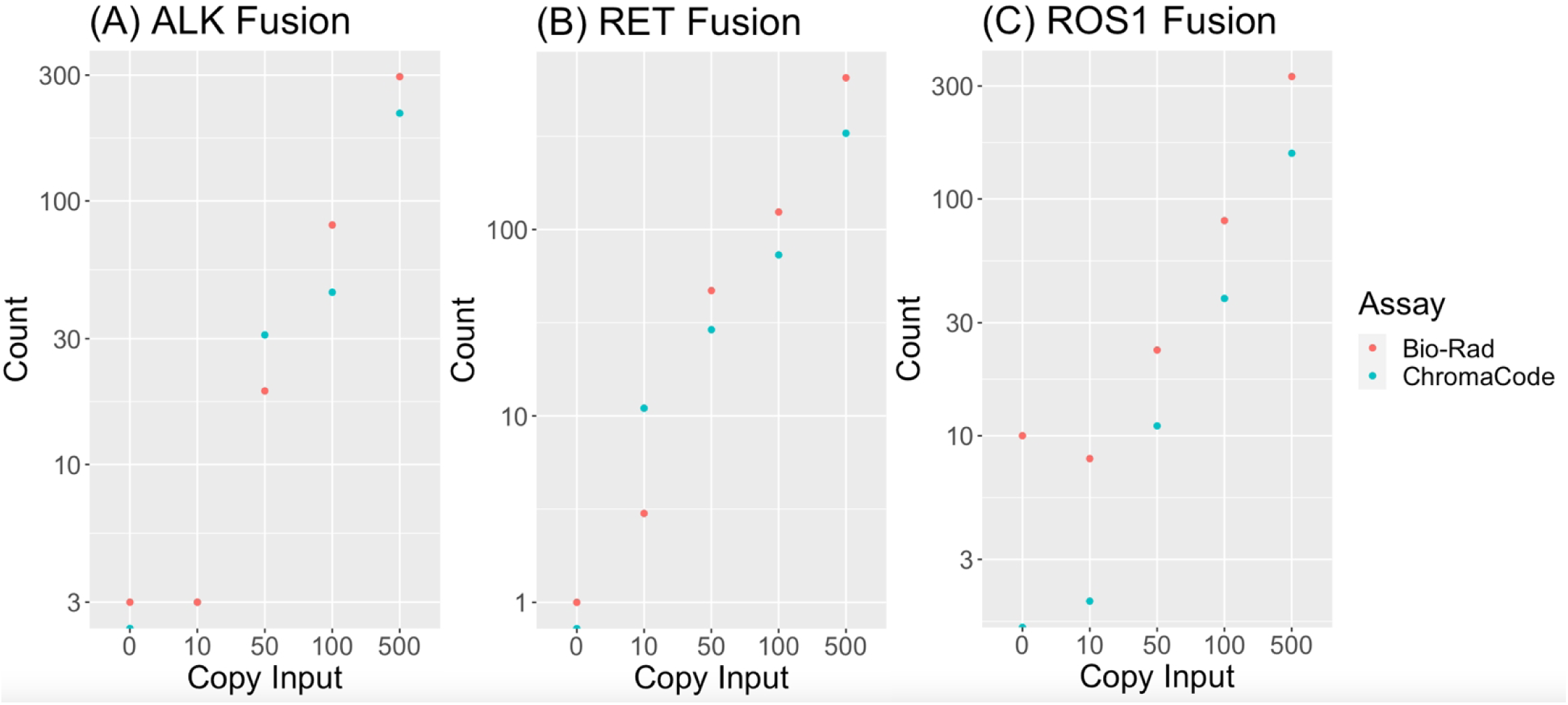
Synthetic titration data for *RET*, *ALK*, *ROS1* with amplitude modulation chemistry and commercially available kit chemistry (BioRad).

**Table S1.**
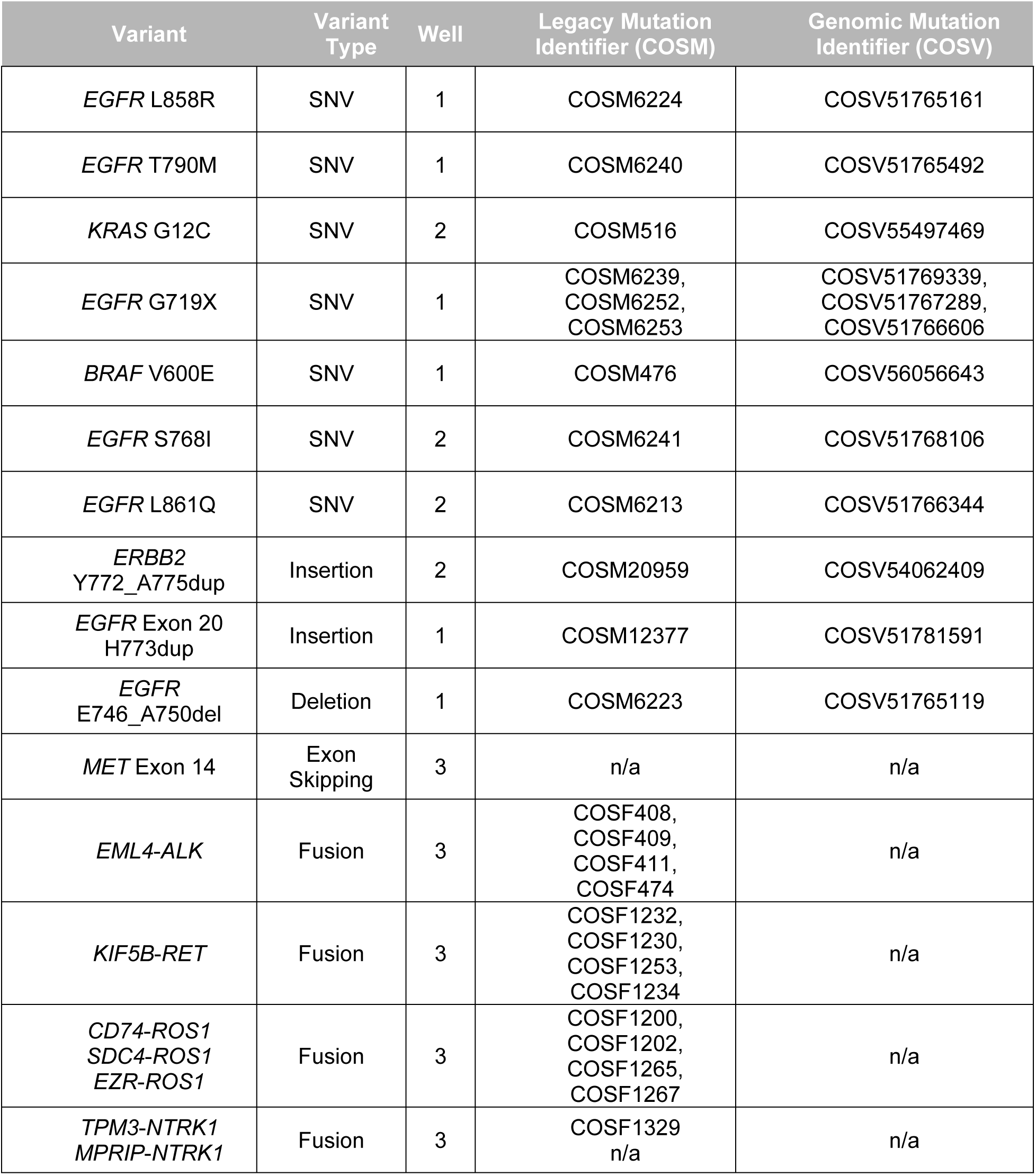
Assay layout for the three well NSCLC dPCR assay inclusive for 12 SNV and indel variants, 14 fusion variants, and *MET* exon skipping.

**Table S1A.**
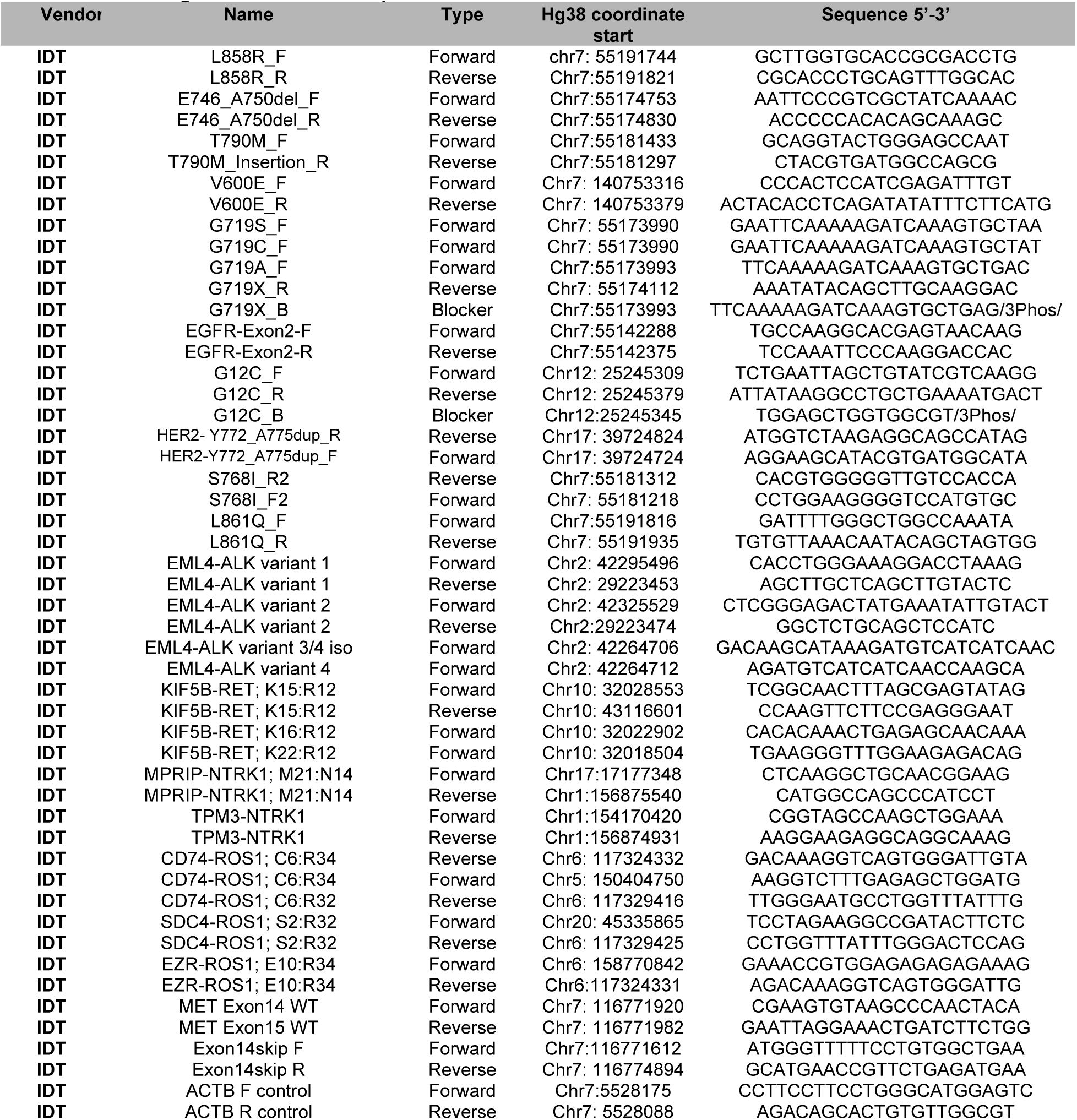
Oligonucleotide sequences

**Table S1B.**
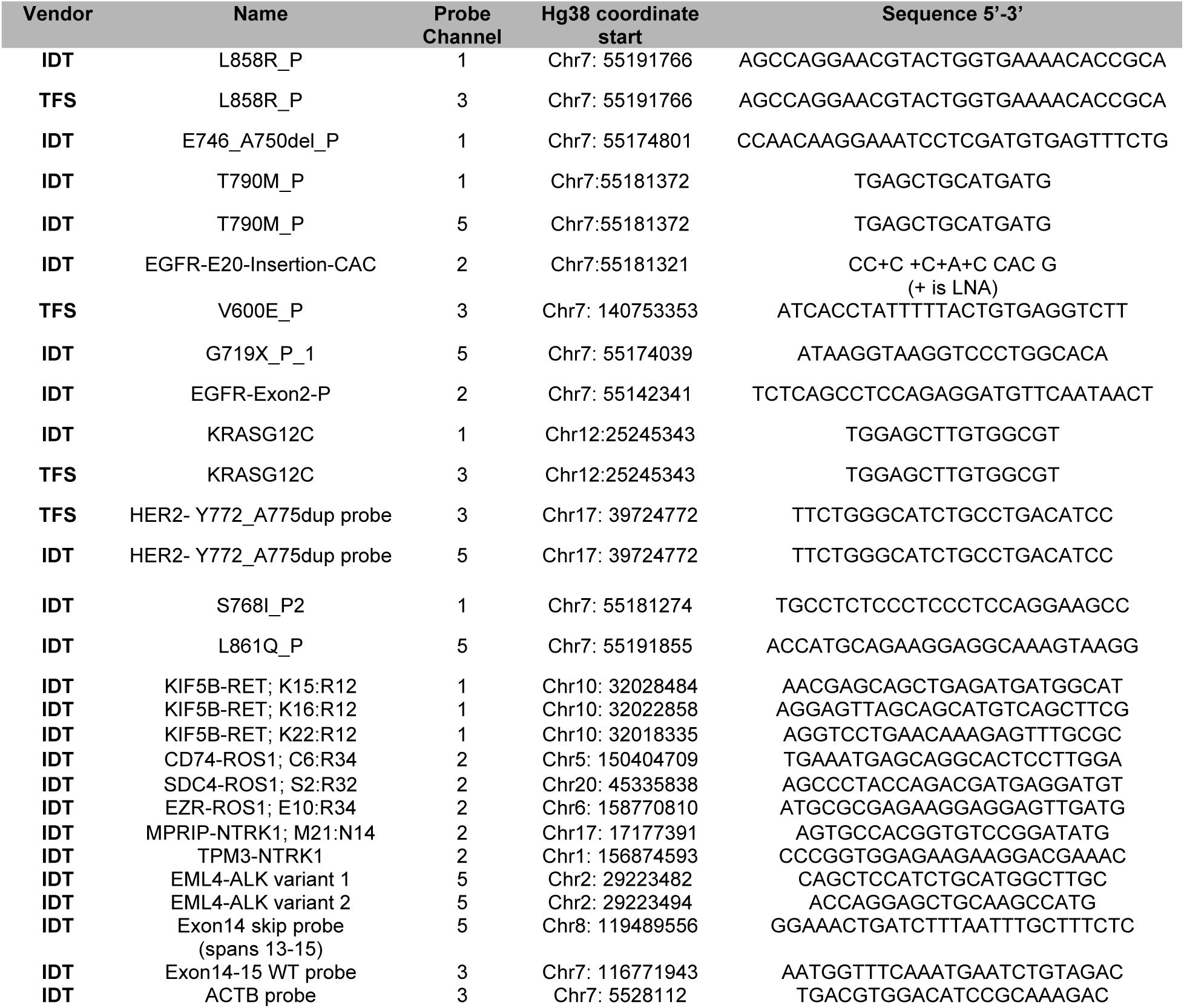
Assay probe sequences.

**Table S2:**
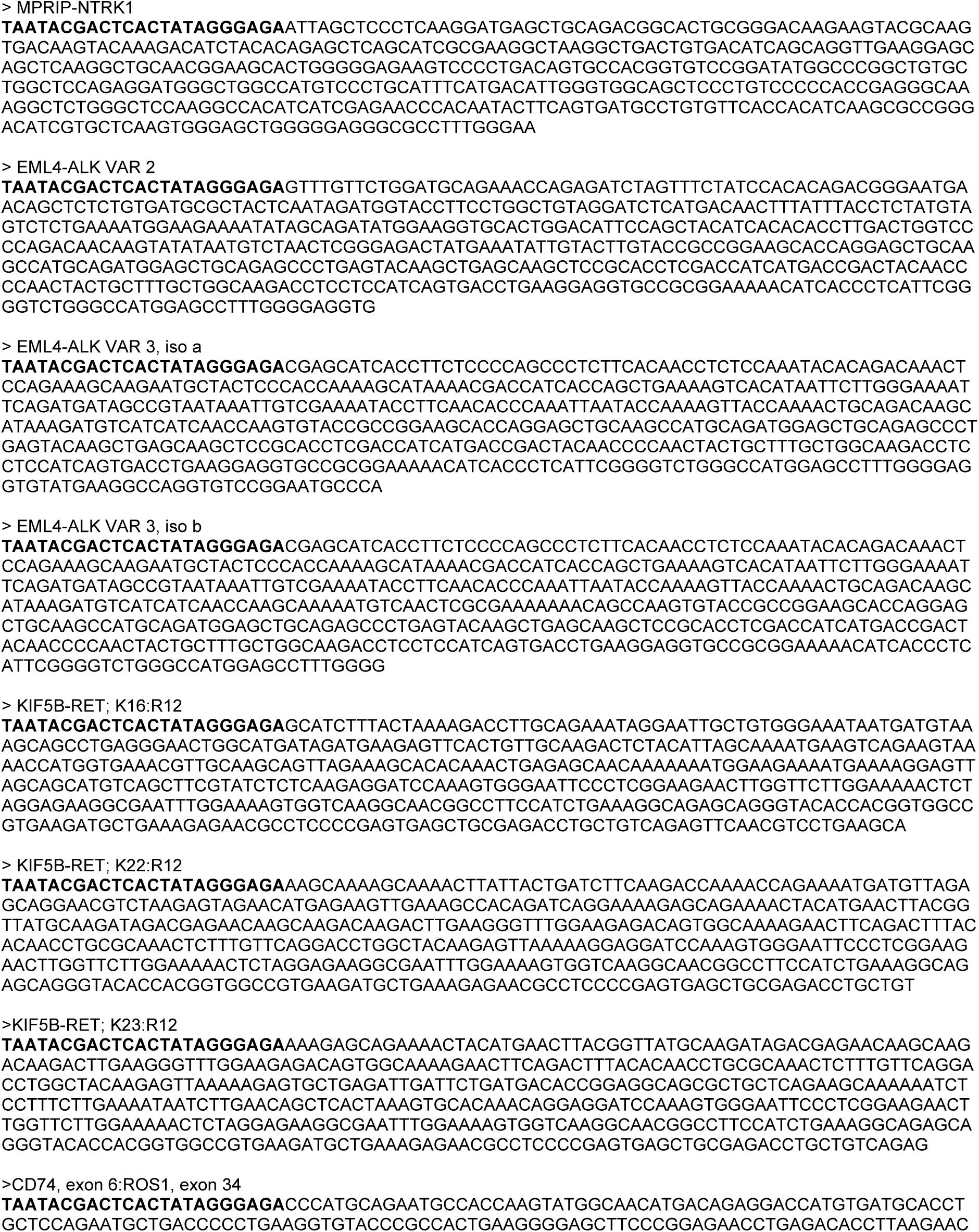
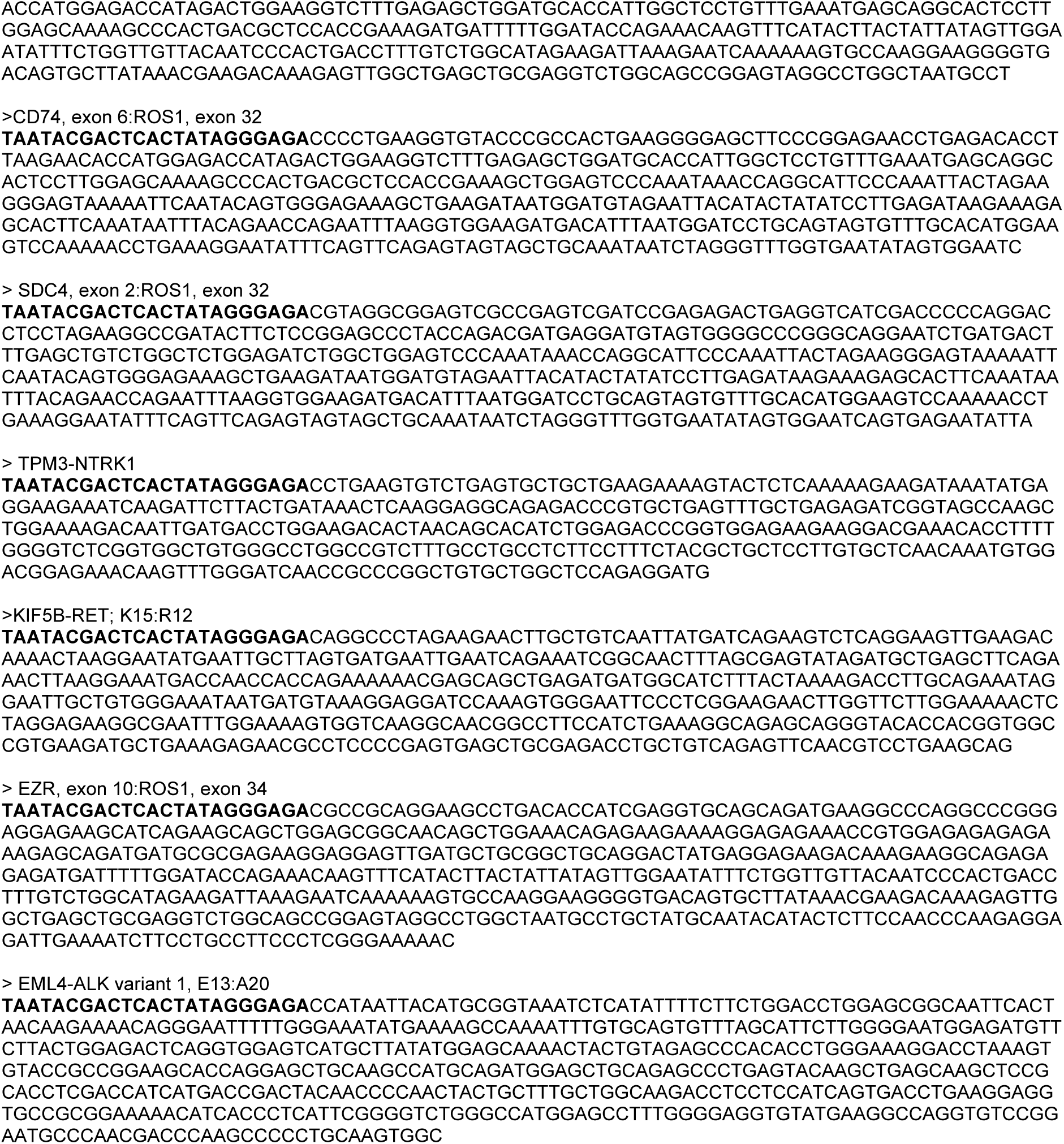
Fusion gBlock Sequences for in vitro transcription of synthetic RNA fusion molecules.

**Table S3.** Oligonucleotide primer-probe composition for amplitude modulation assay. Component name suffixes identify the type of oligo used (‘_F’ = forward primer; ‘_R’ = reverse primer; ‘_B’ = blocker and ‘_P’ = probe).

| Component | Final concentration in dPCR reaction (nM) |
| --- | --- |
| L858R_F | 200 |
| L858R_R | 200 |
| E746_A750del_F | 200 |
| E746_A750del_R | 200 |
| T790M_F | 200 |
| T790M_Insertion_R | 200 |
| V600E_F | 200 |
| V600E_R | 200 |
| G719S_F | 200 |
| G719C_F | 200 |
| G719A_F | 200 |
| G719X_R | 200 |
| G719X_B | 200 |
| EGFR-Exon2-F | 200 |
| EGFR-Exon2-R | 200 |
| L858R_P | 22.4 |
| L858R_P | 20 |
| E746_A750del_P | 20 |
| T790M_P | 18.3 |
| T790M_P | 20 |
| EGFR-E20-Insertion-CAC_P | 20 |
| V600E_P | 5.8 |
| G719X_P_1 | 6.7 |
| EGFR-Exon2-P | 8.5 |

DNA Well #2
| Component | Final concentration in dPCR reaction (nM) |
| --- | --- |
| EGFR-Exon2-F | 300 |
| EGFR-Exon2-R | 300 |
| G12C_F | 300 |
| G12C_R | 300 |
| G12C_B | 300 |
| HER2- Y772_A775dup_R | 300 |
| HER2-Y772_A775dup_F | 300 |
| S768I_R2 | 300 |
| S768I_F2 | 300 |
| L861Q_F | 300 |
| L861Q_R | 300 |
| EGFR-Exon2-P | 20 |
| KRASG12C_P | 18 |
| KRASG12C_P | 18.6 |
| HER2- Y772_A775dup probe | 20 |
| HER2- Y772_A775dup probe | 20 |
| S768I_P2 | 20 |
| L861Q_P | 12 |

RNA Well #3
| Component | Final Concentration in dPCR Reaction (nM) |
| --- | --- |
| EML4-ALK variant 1 F | 300 |
| EML4-ALK variant 1 R | 900 |
| EML4-ALK variant 2 F | 300 |
| EML4-ALK variant 2 R | 300 |
| EML4-ALK variant 3/4 iso F2 | 300 |
| EML4-ALK variant 4 F | 600 |
| KIF5B-RET; K15:R12 (F2) | 300 |
| KIF5B-RET; K15:R12 (R1) | 1200 |
| KIF5B-RET; K16:R12 (F2) | 300 |
| KIF5B-RET; K22:R12 (F2) | 600 |
| MPRIP (ex 21) - NTRK1 (ex 14) (F2) | 300 |
| MPRIP (ex 21) - NTRK1 (ex 14) (R2) | 300 |
| TPM3-NTRK1 (F2) | 300 |
| TPM3-NTRK1 (R2) | 300 |
| CD74, ex6:ROS1, ex34 (R2) | 300 |
| CD74, ex6:ROS1, ex34 (F1) | 600 |
| CD74, ex6:ROS1, ex32 (R1) | 300 |
| SDC4, ex2:ROS1, ex32 (F3) | 300 |
| SDC4, ex2:ROS1, ex32 (R3) | 300 |
| EZR, ex10:ROS1, ex34 (F1) | 300 |
| EZR, ex10:ROS1, ex34 (R1) | 300 |
| Exon14 WT F | 300 |
| Exon15 WT R | 300 |
| Exon14skip F (exon13) | 300 |
| Exon14skip R (exon15) | 300 |
| ACTB F control | 300 |
| ACTB R control | 300 |
| KIF5B-RET; K15:R12 (P2) | 144 |
| KIF5B-RET; K16:R12 (P2) | 107 |
| KIF5B-RET; K22:R12 (P1) | 75 |
| KIF5B-RET; K22:R12 (P1) | 125 |
| CD74, ex6:ROS1, ex34 (P1) | 95 |
| CD74, ex6:ROS1, ex34 (P1) | 75 |
| SDC4, ex2:ROS1, ex32 (P3) | 116 |
| EZR, ex10:ROS1, ex34 (P1) | 101 |
| MPRIP (ex 21) - NTRK1 (ex 14) (P1) | 12 |
| TPM3-NTRK1 (P1) | 13 |
| EML4-ALK variant 1 P | 84 |
| EML4-ALK variant 1 P | 75 |
| EML4-ALK variant 1 P | 120 |
| EML4-ALK variant 2 P | 96 |
| Exon14skip probe (spans 13-15) | 29 |
| Exon14-15 WT probe | 392 |
| ACTB probe | 90 |

**Table S4.** Design of contrived sample experiments. Each reportable in Table 1 was tested one at a time by combining wild-type human biological genomic DNA and synthetic oligonucleotides containing each variant of interest. In addition, combinations of the most common variant combinations (*EGFR* L858R + *EGFR* T790M, *EGFR* T790M + *EGFR* Exon 19 del, *EGFR* Exon 19 del + *EGFR* L858R, and all three of these together) were tested under the same copy number conditions.

| DNA Background Type | Total mass of background DNA (from Absolute Q) | Copy number of synthetic variant DNA (gBlock) | Replicates at each copy number |
| --- | --- | --- | --- |
| FFPE DNA | 21 ng | 2320 | 3 |
|  |  | 1160 | 3 |
|  |  | 464 | 3 |
|  |  | 232 | 4 |
|  |  | 116 | 4 |
|  |  | 58 | 4 |

| RNA Background Type | ACTB copy number of background RNA (from Absolute Q) | Copy number of RNA Fusion (IVT product) | Replicates at each copy number |
| --- | --- | --- | --- |
| FFPE RNA | 5000 | 5000 | 3-10 |
|  |  | 7500 | 3-10 |
|  |  | 10000 | 3-10 |
|  |  | 11250 | 10 |

**Table S5.** The amplitude modulation assay for RNA fusions was benchmarked using a commercially available fusion RNA reference mix (SeraSeq Fusion RNA Mix v4, 0710-0497, SeraCare) (number of replicates = 4). The reference and the assay shared inclusivity for CD74-ROS1, EML4-ALK var 1, and MET Exon 14 skipping; they did not share inclusivity for RET, NTRK1, ACTB, or MET wild type.

| Channel | Variant | Amplitude modulation measured concentration (cp/uL) | Certificate of Analysis concentration (cp/uL) |
| --- | --- | --- | --- |
| Channel 2 | CD74-ROS1;<br>C6:R34 | 104.5 | 113.4 |
| Channel 5 | MET Ex 14 Skip | 153.6 | 125.4 |
| Channel 5 | EML4-ALK, var 1 | 57.6 | 86.5 |

**Table S6a.**
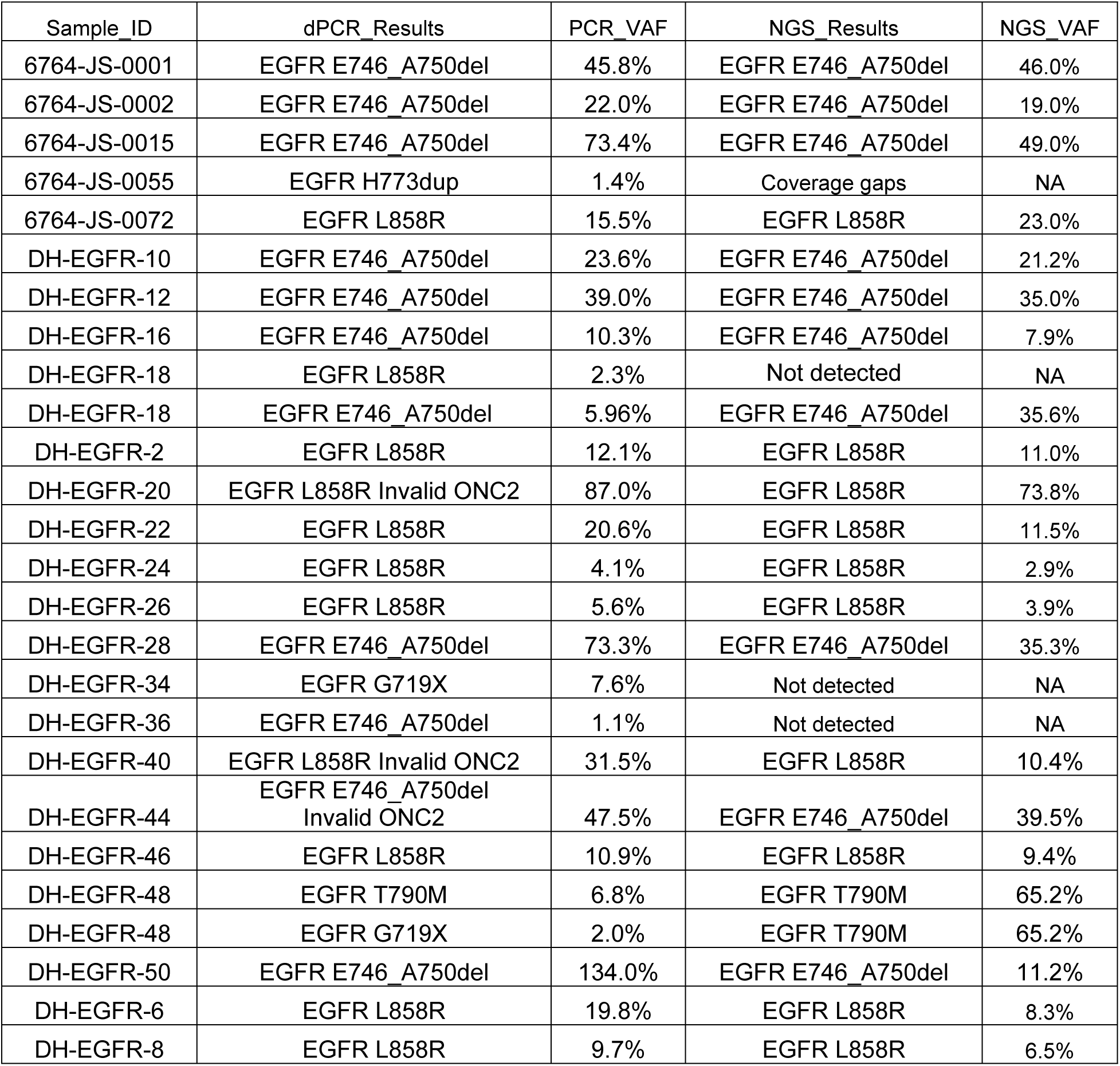
Clinical sample results comparing dPCR to NGS. All samples passed NGS QC and were positive in either NGS or dPCR.

**Table S6b.**
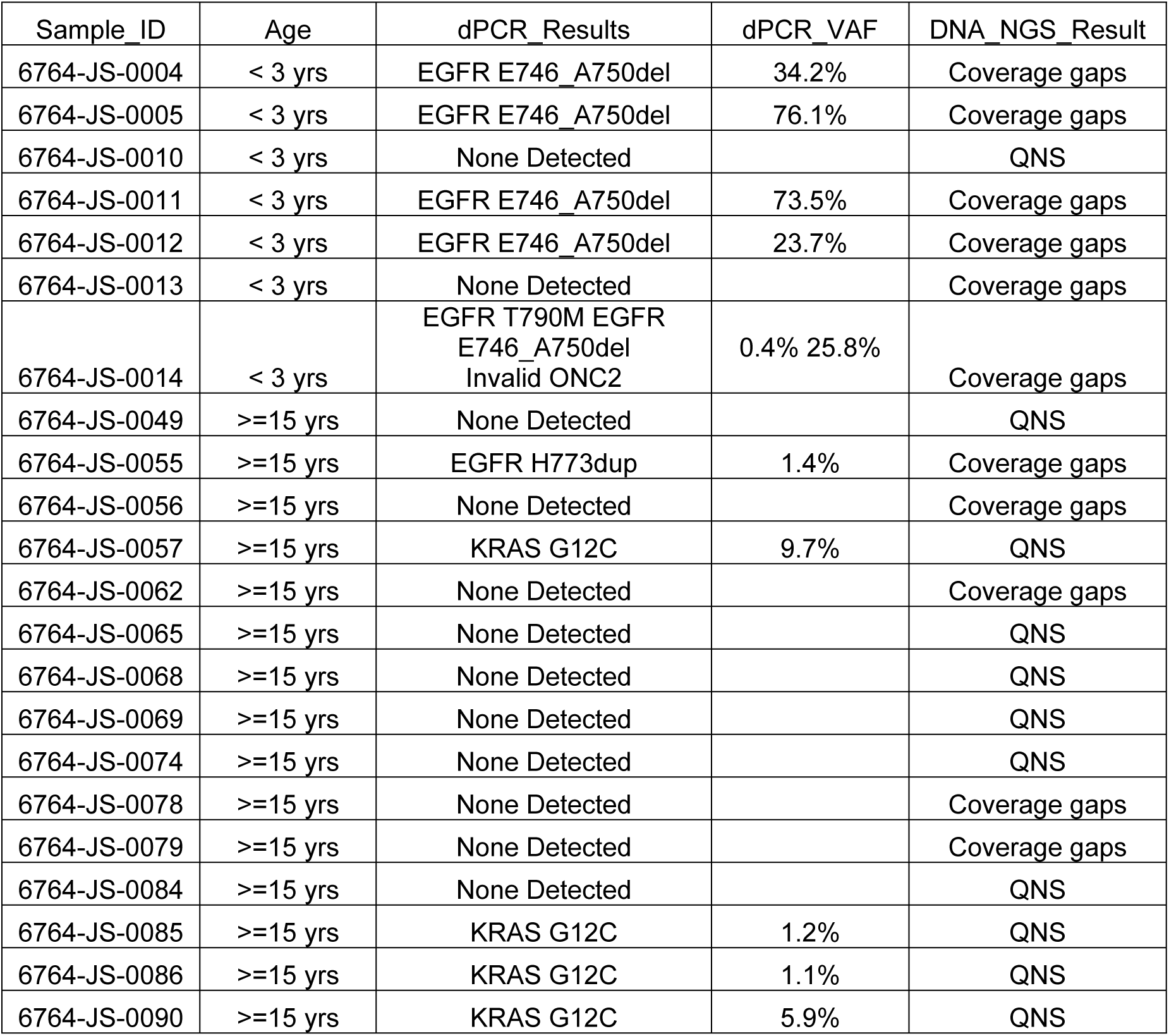
Clinical dPCR results for FFPE samples that either failed to sequence or had gaps in sequence coverage, while the dPCR assay generated valid results.

| Sample_ID | Age | dPCR_Results | dPCR_VAF | DNA_NGS_Result |
| --- | --- | --- | --- | --- |
| 6764-JS-0004 | < 3 yrs | EGFR E746_A750del | 34.2% | Coverage gaps |
| 6764-JS-0005 | < 3 yrs | EGFR E746_A750del | 76.1% | Coverage gaps |
| 6764-JS-0010 | < 3 yrs | None Detected |  | QNS |
| 6764-JS-0011 | < 3 yrs | EGFR E746_A750del | 73.5% | Coverage gaps |
| 6764-JS-0012 | < 3 yrs | EGFR E746_A750del | 23.7% | Coverage gaps |
| 6764-JS-0013 | < 3 yrs | None Detected |  | Coverage gaps |
| 6764-JS-0014 | < 3 yrs | EGFR T790M EGFR E746_A750del<br>Invalid ONC2 | 0.4% 25.8% | Coverage gaps |
| 6764-JS-0049 | >=15 yrs | None Detected |  | QNS |
| 6764-JS-0055 | >=15 yrs | EGFR H773dup | 1.4% | Coverage gaps |
| 6764-JS-0056 | >=15 yrs | None Detected |  | Coverage gaps |
| 6764-JS-0057 | >=15 yrs | KRAS G12C | 9.7% | QNS |
| 6764-JS-0062 | >=15 yrs | None Detected |  | Coverage gaps |
| 6764-JS-0065 | >=15 yrs | None Detected |  | QNS |
| 6764-JS-0068 | >=15 yrs | None Detected |  | QNS |
| 6764-JS-0069 | >=15 yrs | None Detected |  | QNS |
| 6764-JS-0074 | >=15 yrs | None Detected |  | QNS |
| 6764-JS-0078 | >=15 yrs | None Detected |  | Coverage gaps |
| 6764-JS-0079 | >=15 yrs | None Detected |  | Coverage gaps |
| 6764-JS-0084 | >=15 yrs | None Detected |  | QNS |
| 6764-JS-0085 | >=15 yrs | KRAS G12C | 1.2% | QNS |
| 6764-JS-0086 | >=15 yrs | KRAS G12C | 1.1% | QNS |
| 6764-JS-0090 | >=15 yrs | KRAS G12C | 5.9% | QNS |

**Extended Table S7: Clinical sample molecular data:**

TableS7_NSCLC_Clinical_Samples.xlsx

**Extended Table S8: Clinical sample metadata:**

TableS8_Clinical_Metadata.xlsx

